# An *In Vitro* Vascularized Micro-Tumor Model of Human Colorectal Cancer Recapitulates *In Vivo* Drug Responses

**DOI:** 10.1101/2020.03.03.973891

**Authors:** Stephanie J. Hachey, Silva Movsesyan, Quy H. Nguyen, Giselle Burton-Sojo, Ani Tankanzyan, Jie Wu, Tuyen Hoang, Michaela M. Hatch, Da Zhao, Elizabeth Celaya, Samantha Gomez, George T. Chen, Ryan T. Davis, Kevin Nee, Nicholas Pervolarakis, Devon A. Lawson, Kai Kessenbrock, Abraham P. Lee, Marian L. Waterman, Christopher C.W. Hughes

## Abstract

Around 95% of anti-cancer drugs that show promise during preclinical study fail to gain FDA-approval for clinical use. This failure of the preclinical pipeline highlights the need for improved, physiologically-relevant *in vitro* models that can better serve as reliable drug-screening tools. The vascularized micro-tumor (VMT) is a novel three-dimensional model system that recapitulates the complex human tumor microenvironment, including perfused vasculature, within a transparent microfluidic device, allowing real-time study of drug responses and tumor-stromal interactions. Here we have validated the VMT platform for the study of colorectal cancer (CRC), the second leading cause of cancer-related deaths, by showing that gene expression, tumor heterogeneity, and treatment response in the VMT more closely model CRC tumor clinicopathology than current standard drug screening modalities, including 2-dimensional (2D) monolayer culture and 3-dimensional (3D) spheroids.

## Introduction

Cancer accounts for 25% of US deaths^1^, and less than 5% of drugs entering clinical trials ultimately receive FDA approval^2^ with most anti-cancer agents failing in clinical trials despite showing promise during preclinical studies. The conclusion is that current model systems are poor predictors of drug response in humans. Traditionally, drug candidates are first tested in 2D cell monocultures. However, cell growth in 2D versus 3D environments not only promotes phenotypic changes in cell morphology, response to stimuli, cell functions and gene expression patterns, but also alters response to therapeutic agents^3–7^. Indeed, both normal and cancerous cells maintain their specific functions in the body owing to the 3D conformation they adopt, encompassing heterogeneous and dynamic cell-cell and cell-matrix interactions^8–10^. Prior efforts to develop appropriate 3D tumor tissues have been limited to the generation of avascular spheroids in artificial matrix, which fail to recapitulate the structure of a vascularized tumor mass, are not viable for long-term study^11,12^, and perhaps most importantly cannot be generated from invasive and often metastatic cell lines that have reduced expression of adhesion molecules. Although animal models have significantly advanced our understanding of complex diseases such as cancer, and are important in the drug development pipeline, these same models require substantial time and resources and species-specific differences can impede clinical translation of results^13,14^.

To address the need for improved preclinical models, we have designed, fabricated and validated a microfluidic device that supports the formation of a perfused, vascularized micro-tumor (VMT) via co-culture of multiple cell types in an extracellular matrix^15–19^. Importantly, growth of the tumor and delivery of therapeutics to the tumor is entirely dependent on “blood” flow through the living vascular network. Physiologic flow rates through the microvasculature are maintained by a gravity-driven pressure differential that does not require external pumps or valves^20^. The VMT (aka ‘tumor-on-a-chip’) is composed entirely of human cells, is optically compatible for real-time fluorescent image analysis and is arrayed for high throughput experiments^21^. Drugs can be tested within days on limited numbers of cells expanded to form the VMTs, providing high sensitivity and rapid turnaround of results.

CRC represents a major source of morbidity and mortality as the second leading cause of cancer-related deaths in the US and the third most common cancer worldwide, with over a million diagnoses and half a million deaths each year. The majority (75%) of patients present with localized disease (stage I and II) and undergo intentionally curative surgery. However, half of these patients will develop, and ultimately die from advanced disease with a 5-year survival rate for stage III and IV disease at only 10%, highlighting the need for more effective therapies. Achieving this will require more accurately modeling tumor biology *in vitro*. Here we show that when CRC cell gene expression, heterogeneity, growth and response to standard chemotherapy are examined, cells grown in the VMT much more closely resemble those grown as xenografts than do cells grown either as monolayers (2D) or as spheroids (3D). Our studies reveal that the VMT recapitulates key features known to drive CRC disease progression and therapeutic failure, including contributions from the microenvironment, while also capturing unique cellular populations and expression signatures not observable in simple *in vitro* models. Specifically, we show how changes in gene expression within the tumor and stroma may underpin drug sensitivities. Our findings support the VMT model as a powerful tool for drug development and disease modeling in oncology.

## Results

### The VMT serves as a platform for improved modeling and direct visualization of the complex tumor microenvironment

We have previously shown the VMT model to be a robust system for disease modeling and drug screening studies^18,19,21^. Multiple tissue units are incorporated within a single microfluidic device that is fitted onto a bottom-less 96-well plate to allow each VMT to be independently treated (Figure 1A). In response to physiologic flow driven by a hydrostatic pressure gradient across the tissue, endothelial cells (EC), fibroblasts and cancer cells introduced into each tissue unit self-organize within an extracellular matrix to form a complex tumor micro-ecosystem by day 5 of VMT culture (Figure 1B). Importantly, tumor growth is entirely dependent on delivery of nutrients through the living vascular network. COMSOL Multiphysics simulation on a fully formed, anastomosed and perfused vascular network shows that the surface velocity of medium flowing through the vessels varies across the tissue, with some areas experiencing higher flow than others, mimicking blood flow through a capillary network *in vivo* (Figure 1C).

**Figure 1:**
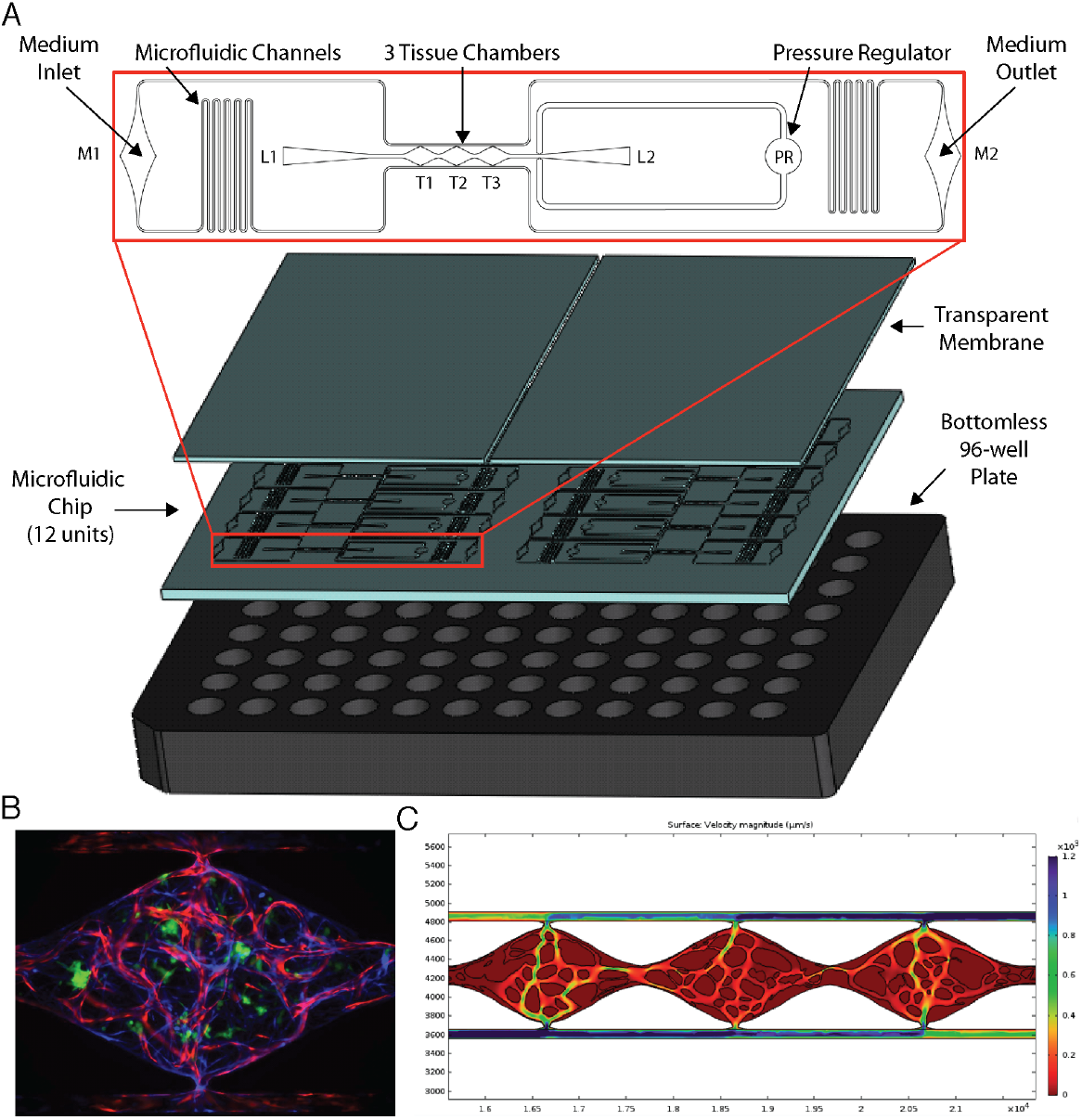
Vascularized micro-tumors (VMTs) are supported by gravity-driven flow through living human microvessels within a microfluidic platform arrayed for high throughput experiments. (A) The microfluidic chip is bonded to a bottom-less 96-well plate via chemical glue and oxygen plasma. Zoomed view shows a single device unit with 3 tissue chambers (T1-3) fed through microfluidic channels, 2 loading ports (L1-2), medium inlet and outlet (M1-2), and a pressure regulator (PR) to prevent gel bursting during loading. (B) Fluorescent image shows a vascularized micro-tumor with EC (mCherry, red) forming the vascular network, fibroblasts (Az, blue) supporting the tissue, and HCT116 cancer cells (GFP, green). Tissue chamber is ~2 × 1 × .1 mm. (C) COMSOL simulation on a perfused vascular network shows the surface velocity of medium flowing through the vessels.

Immunofluorescence staining reveals stark differences in collagen density and organization between the VMO and VMT. Collagen III in particular is highly expressed in the VMT, especially near tumor cells, and forms fiber ‘tracks’ throughout the tissue. Expression is more diffuse in the VMO (Supplemental Figure 1A–C). Intriguingly, live imaging studies have shown that cancer cells migrate rapidly on collagen fiber tracks in areas enriched in collagen^22^. To capture the full sequence of events that comprise the coordinated self-organization of the VMT as it progresses from single cancer cells embedded in a stromal tissue niche to a fully-formed vascularized tumor mass, we performed a 10-day time-lapse study (Supplemental Videos 7, 8). We observed a remarkable series of events: tumor cells divide slowly while the vasculature forms, but once formed and flow is established the microtumors grow rapidly. Remarkably, some of these microtumors undergo a rapid and explosive disaggregation into single cells that migrate radially from the origin, suggesting a coordinated EMT event (Supplemental Figure 1D). Taken together, the defining features of the VMT are changes in ECM composition associated with *in vivo* tumor progression, and the presence of an abnormal vasculature that closely models known characteristics of tumor-associated vasculature *in vivo* (see below).

### Vessels in the VMT are irregular and leaky, hallmarks of *in vivo* tumors

Distinct changes in tumor-associated vascular architecture and function are known to occur during tumor progression. Although tumors recruit blood vessels to support their growth, the resulting vasculature is irregular, leaky, and ill-perfused, likely as a result of disrupted microenvironmental cues such as growth factor availability and matrix composition^23–25^. Comparisons between vessels in the absence of tumor, the vascularized micro-organ (VMO), and the VMT show that the vessels in the VMT recapitulate key features of *in vivo* tumor-associated vasculature, with vessel irregularity and compression becoming more pronounced at later time points, as the tumors grow larger (Figure 2A). These changes are manifest as increases in the number of blind-ended vessels and an overall decrease in vessel length and diameter, and number of junctions (Figure 2B-E). In line with findings *in vivo*, tumor vessels in the VMT also show increased leakiness (Figure 2F, G). Furthermore, as a result of structural heterogeneity in tumor-associated vasculature, the VMT shows differential patterns of perfusion, with highly leaky areas found adjacent to ill- or non-perfused areas (Supplemental Videos 1-6). Overall, the vasculature within the VMT is significantly more leaky than the VMO, indicating that the vessels are affected by the presence of the tumor, as they are *in vivo*.

**Figure 2:**
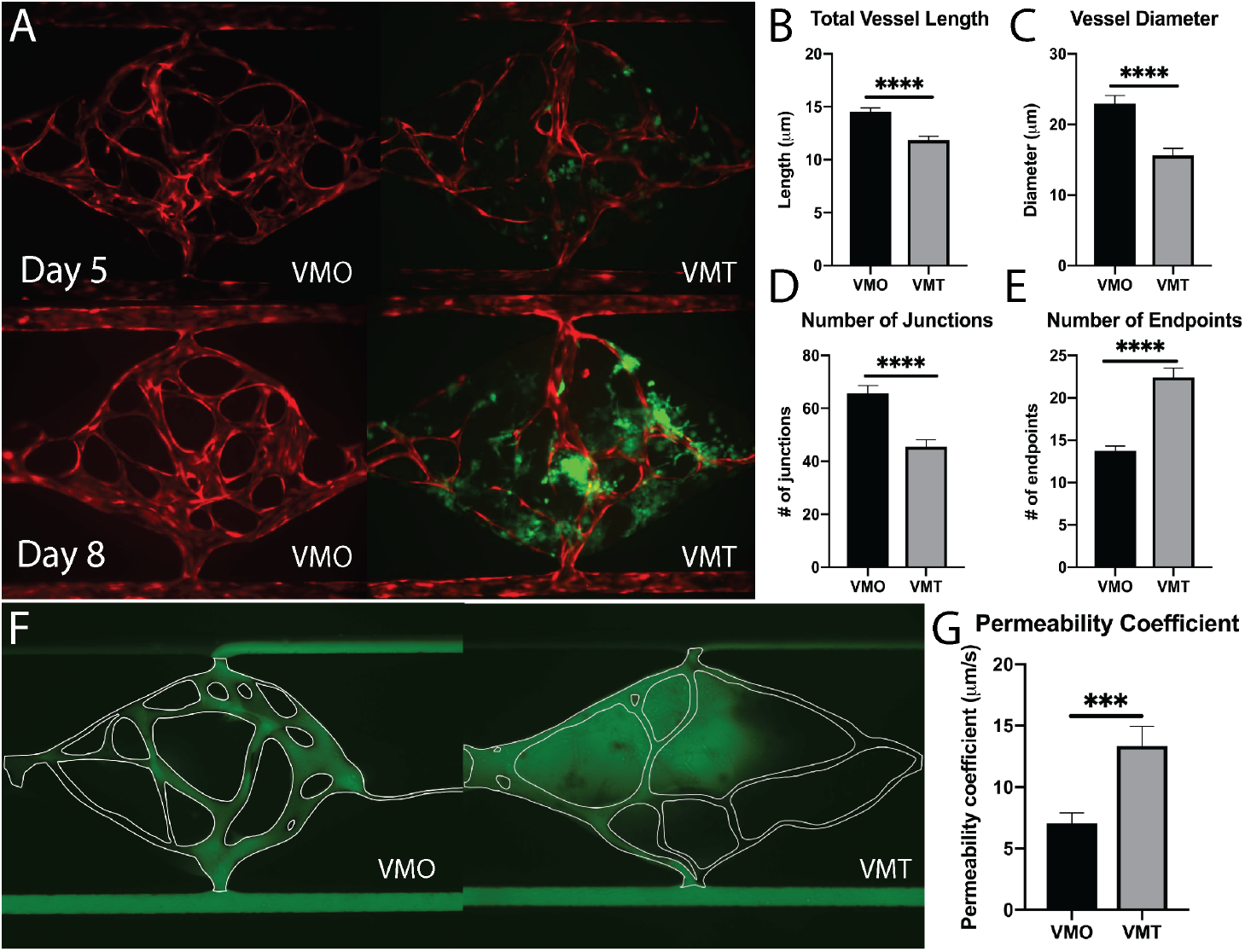
The VMT models key features of *in vivo* tumor vasculature. (A) Comparison of VMO and VMT on days 5 and 8 show irregular vasculature in the VMT that worsens over time. EC (mCherry, red), HCT116 cancer cells (GFP, green). (B) Quantification of total vessel length, (C) vessel diameter, (D) number of junctions and (E) number of endpoints. While vessel length, diameter and density are decreased in the VMT, the number of endpoints increases, suggesting angiogenic sprouting in response to the tumor. (F) Perfusion with 70 kD FITC dextran reveals fully patent networks in the VMO, with leaky and non-perfused vessels in the VMT. (G) Quantification of the permeability coefficient shows that the VMT is twice as leaky as the VMO.

### Tumor cells grown in the VMT closely resemble xenograft tumors as revealed by transcriptomic profiling

To test our hypothesis that the VMT more closely models *in vivo* tumors than monolayer cultures, we carried out transcriptomic analysis of 770 cancer-related genes, from 13 cancer-associated pathways (Nanostring, Inc), on HCT116 colorectal cancer (CRC) cells that had been grown either in the VMT, as xenograft tumors, or in 2D monocultures. HCT116 cells expressing mCherry were isolated from all experimental groups (VMT, monolayer and xenograft) by fluorescence activated cell sorting (FACS). Remarkably, we found that the gene expression patterns of tumor cells derived from the VMT closely matched *in vivo* tumor-derived cells (difference not significant), whereas HCT116 cells grown in monolayer did not (difference significant at p<0.0001, Figure 3A). Interestingly, the variation between samples was highest for monolayer cultures (12.8%) and second highest for xenograft tumors (6.6%), with the VMT yielding the least variability between replicates (3.5%), highlighting the robustness and reproducibility of the VMT model. Pathways consistently enriched in the VMT and xenograft-derived CRC cells, but not in 2D monocultures, included PI3K-Akt signaling, MAPK signaling, Ras signaling, FGF signaling, chromosome and micro satellite instability, and epithelial-to-mesenchymal transition (Figure 3B). Interestingly, genes that were consistently differentially expressed in the VMT compared to xenograft were enriched for beta-1 integrin signaling (COL1A1, COL3A1, FN1) and urokinase-type plasminogen activator receptor-mediated signaling (PDGFRB, HGF, MMP3), indicating that cell-matrix interactions may be more active within the VMT.

**Figure 3:**
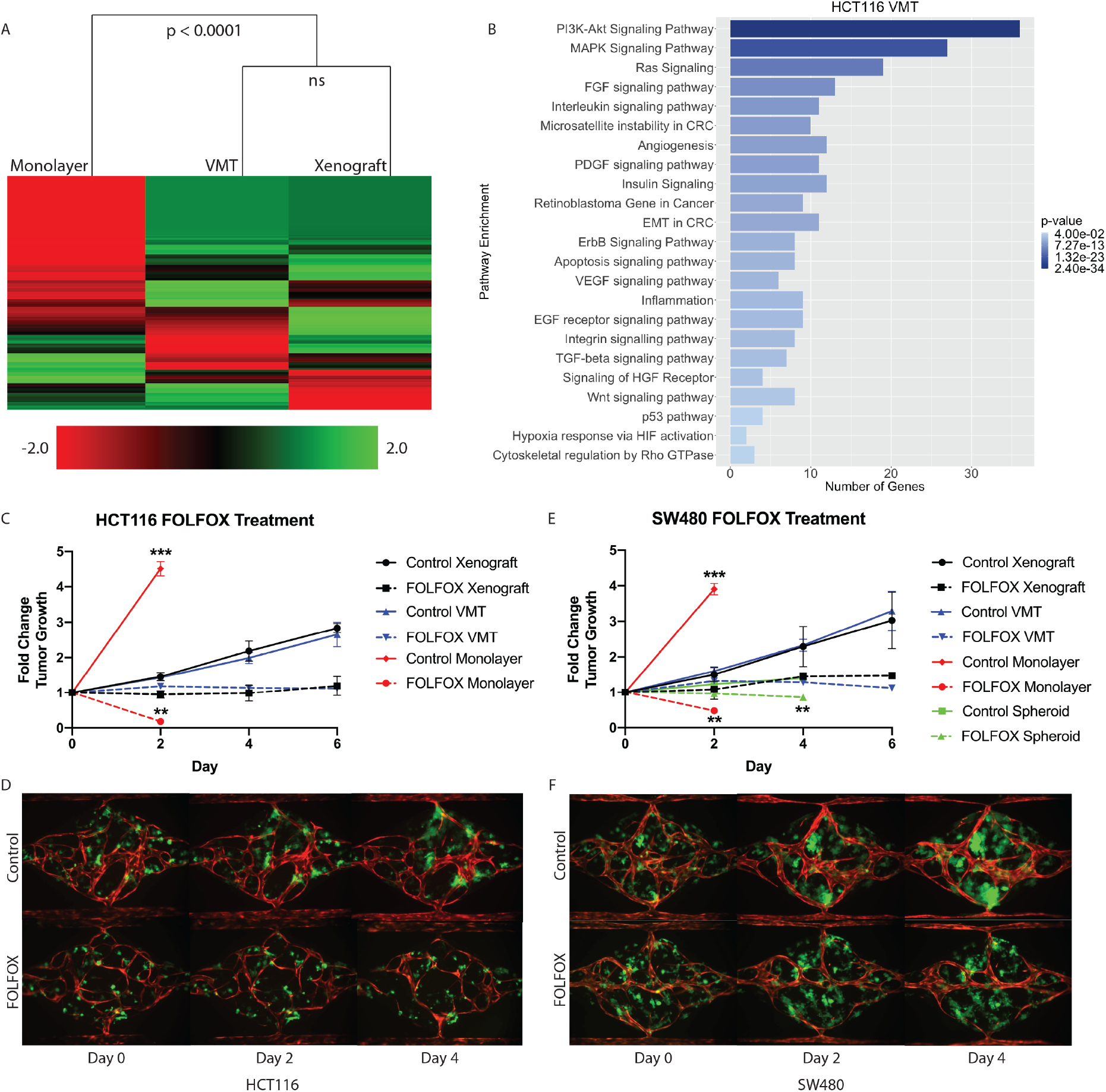
Comparison of gene expression, tumor growth and response to drug treatment between VMT, 2D and 3D monocultures and xenograft tumors. (A) Bulk RNA sequencing of 770 cancer-related genes in HCT116 CRC cell line-derived monolayers, VMT and xenograft tumors. (B) Pathway enrichment plot for VMT similarity to xenograft tumor. (C) Growth curves of HCT116 cells grown in monolayer, in mice or in the VMT and treated with standard of care, FOLFOX. (D) Representative images of HCT116-VMT. (E) Growth curves of SW480 cells in monolayer, spheroids, VMT and mice treated with FOLFOX. (F) Representative images of SW480-VMT. ** p < 0.001, *** p < 0.0001.

### The VMT recapitulates xenograft tumor growth rates and response to standard-of-care therapy

Based on the significant overlap in gene expression between HCT116 cells grown in the VMT platform or as xenografts, we next compared growth rates and drug responses between tumor cells grown in monolayer cultures, as VMTs and as xenograft tumors. To capture the molecular landscape of CRC, we tested two established CRC cell lines originating from tumors of different subtypes: HCT116, which is a primary-derived, MSI-high (microsatellite instable-high), KRAS-mutant CRC line, representing about 15% of clinical CRC cases; as well as SW480, a primary-derived, MSS (microsatellite stable), APC-mutant line representing the majority (85%) of clinical CRC cases. First-line treatment for advanced metastatic CRC is systemic cytotoxic chemotherapy using FOLFOX (a regimen of 5 fluorouracil (5FU), leucovorin and oxaliplatin)^26^. Therefore, we treated each system with pharmaceutical-grade FOLFOX at a dose, duration and frequency that approximated the clinical administration, plasma Cmax and clearance for each drug (see Materials and Methods). Drugs were added directly to monolayer cultures, delivered through the vasculature in VMTs and injected i.p. in mice.

Both HCT116 and SW480 grew rapidly in the 2D culture (and outgrew the well in a couple of days), and were also extremely sensitive to drug treatment. In sharp contrast, the growth of both HCT116 and SW480 was much slower in the VMT format, and drugs retarded tumor growth but did not reverse it. Remarkably, these kinetics completely recapitulated the responses seen when the cells were grown as xenografts (Figure 3C–F). Specifically, HCT116-derived xenograft and VMT tumors showed no statistically significant difference in growth over the entire 6-day experimental period (day 6 fold change from baseline: 2.84±1.31 vs 2.66±0.35, respectively) (Figure 3C, D). In contrast, monolayer cultures exceeded the maximum growth observed in VMT and xenograft tumors in only 48 hours, to reach a fold change of 4.51±0.21. The 2D cultures were significantly different from both VMT and xenograft at the 2-day time point (p-value <0.0001) (Figure 3C). For FOLFOX treatment, we observed that xenograft tumors and VMT showed similar drug sensitivity (1.19±0.27 and 1.12±0.01 fold change, respectively) and thus similar effect size (approximately 58%) at 6 days post-treatment (Figure 3C). Monolayer cultures, however, showed an approximate 96% reduction in tumor growth at the 48-hour time point, which is significantly different from both xenograft and VMT responses (p-value <0.001) (Figure 3C).

We found very similar results with SW480-derived xenograft and VMT tumors, with respective fold change 3.03±0.79 and 3.29±0.56 above baseline on day 6, and no significant difference in growth between VMT and xenograft tumor at any time point (Figure 3E, F). FOLFOX treated xenograft and VMT-derived tumors also showed similar drug sensitivity (Figure 3E), while monolayer cultures showed significantly increased cell growth and drug sensitivity at the 48-hour time point compared to xenograft and VMT (Figure 3E). We also studied SW480 spheroids, and intriguingly, these showed significantly reduced growth compared to xenograft tumors (fold changes at day 4: 1.41±0.06 and 2.29±0.53, respectively) while maintaining roughly the same effect size in response to FOLFOX treatment. However, the absolute response to FOLFOX was significantly different in the spheroid cultures compared to xenograft (0.85±0.16 and 1.45±0.02 fold change in tumor growth, respectively). Hence, for two different subtypes of CRC, we show that while 2D and 3D monocultures are poorly predictive of tumor growth and response to chemotherapy *in vivo*, the VMT accurately predicts both.

### HCT116-derived VMTs retain *in vivo* tumor heterogeneity not seen in 2D monocultures

Based on stark differences in tumor growth and response to chemotherapy between 2D and 3D monocultures and the VMT, we next sought to explore the distinct gene expression changes that occur at the single-cell level when cells are co-cultured within the dynamic 3D tumor microenvironment of the VMT. Here we used single-cell mRNA sequencing (scRNAseq) to profile the transcriptomes of cells isolated from HCT116-containing VMT and matched HCT116 CRC cells, EC and fibroblasts growing in 2D monocultures (Figure 4A). Unbiased clustering analysis using the Seurat pipeline^27^ revealed distinct shifts in gene expression between the cells grown in the VMT and those grown in monolayer (Supplemental Figure 2A,B). Guided by annotated lineage-specific markers in humans^28^, along with prior knowledge, two populations were identified as EC (EC A and EC B) by differential expression of PECAM1 (CD31+), two populations were derived from fibroblasts (fibroblast A and B) based on high expression of MMP2 and ACTA2, two populations were enriched for EPCAM and labeled as tumor (A and B), and a third tumor cluster highly expressed KRT5 (tumor C) (Supplemental Figure 2C,D). Overall, the cells derived from the VMT showed higher expression of cell-type specific markers than the same cells grown in 2D cultures (Figure 4B), including the epithelial marker EPCAM. Intriguingly, two populations of cells were found in the VMT but not in 2D monocultures (Figure 4C). The first of these showed higher KRT5 expression, characteristic of cells undergoing epithelial-to-mesenchymal transition (EMT)^29^ (Supplemental Figure 2E, F) and this finding was confirmed by immunofluorescence staining (Figure 4D).

**Figure 4:**
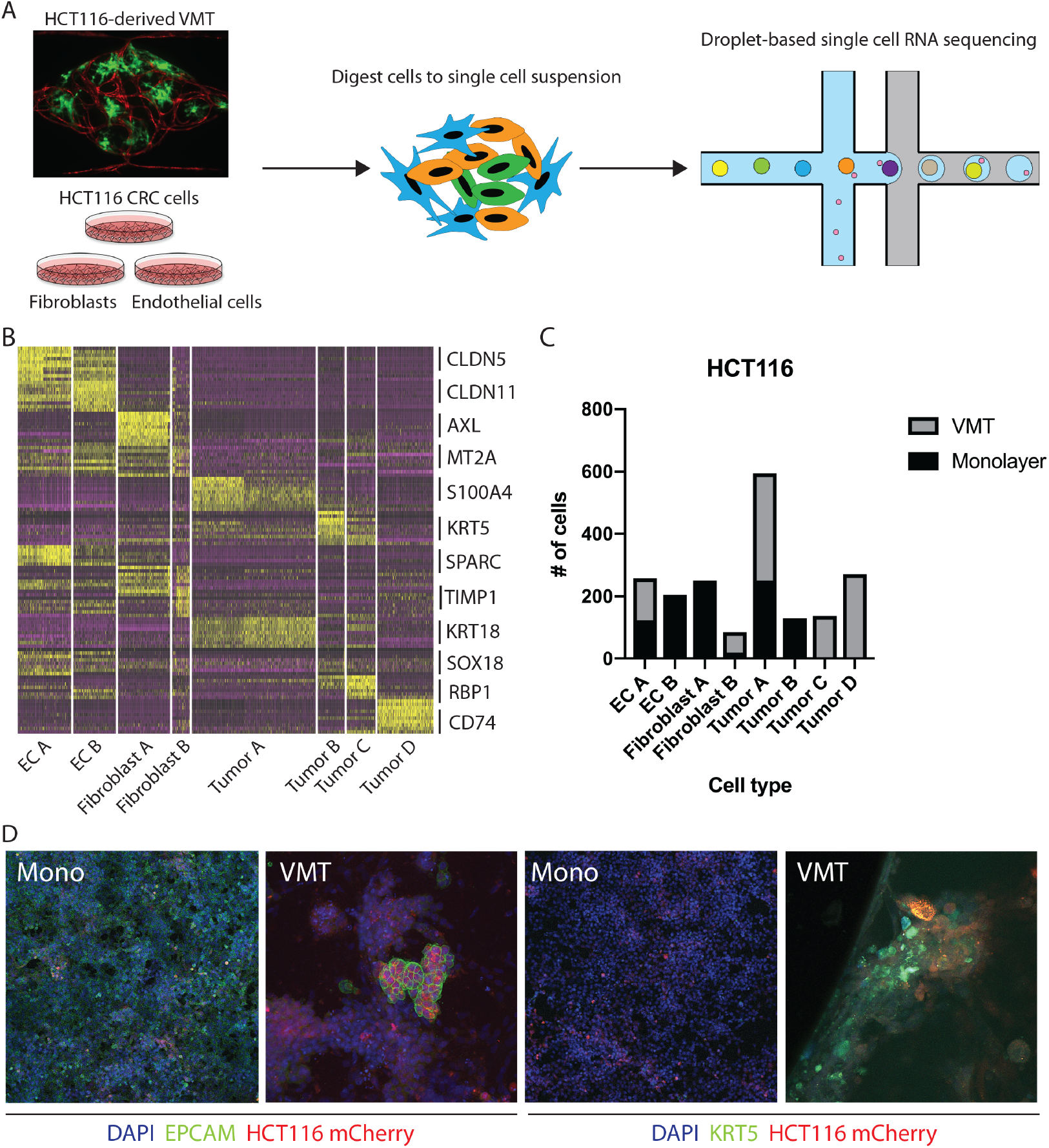
scRNAseq reveals HCT116 tumor heterogeneity in the VMT platform. (A) Experimental design. HCT116 CRC cells, EC and fibroblasts were grown individually as 2D monocultures or co-cultured within the VMT, harvested on day 7 of culture and processed for droplet-based single cell RNA sequencing. (B) Heatmap of top 10 differentially-expressed genes. (C) Distribution of cell types by sample. (D) Immunofluorescent staining of EPCAM and KRT5.

The second cluster of cells unique to the VMT showed differential expression of CSPG4 (NG2+, a pericyte marker), however, additional pericyte markers such as PDGFRβ and DES were absent (Supplemental Figure 3A). Furthermore, expression of PAGE5 (a proposed tumor marker) and VGF (a neuroendocrine marker) were restricted to this population, hinting at a tumor origin, although other markers expressed in highly invasive and resistant CRC tumors, such as CHGA and SYP, were absent^30,31^.

Single nucleotide variation (SNV) profiling of raw reads (see Materials and Methods) allowed us to identify the cells as HCT116 in origin, but with enrichment of CNV on chromosomes 1, 11 and 19 compared to the other tumor populations. While tumor populations A and B had a similar CNV profile, C clustered with tumor D, and both showed distinguishing patterns of CNV enrichment, suggesting a greater degree of tumor heterogeneity within the VMT, compared to monolayer, likely through selection of HCT116 CRC subclones (Supplemental Figure 3B). Taken together, these analyses revealed a higher degree of heterogeneity within VMT-derived tumor subpopulations relative to monolayer cultures.

In Supplemental Figure 2B, tSNE plotting shows the degree to which subpopulations were unique to either VMT or monolayer culture, and a heat map with the combined top 10 cluster-discriminative genes for HCT116 populations showed that clusters were distinct in their gene signatures (Figure 4B). The distribution of cells from each sample based on cell type reveals distinct changes in the VMT compared to 2D monocultures (Figure 4C). Importantly, we found greater tumor heterogeneity in the VMT compared to monolayer, with tumor A, C and D all being present in the VMT, whereas monolayer cultures were comprised predominantly of tumor type A. The stroma was also distinctly different in the VMT compared to monolayer, with EC B and Fibroblast A populations being diagnostic of monolayer culture. Immunofluorescence (IF) staining of HCT116 in the VMT or growing in a monolayer confirmed the presence of tumor subpopulations in the VMT (Epcam+, Cytokeratin 5+), with only a subset of the tumors expressing EPCAM in the VMT, and a subset of cells expressing high levels of KRT5. In contrast, HCT116 cells growing in 2D showed ubiquitous but low expression of EPCAM and low levels of KRT5 in a subpopulation of cells (Figure 4D).

### Lineage hierarchy reconstruction reveals a unique VMT-derived HCT116 tumor population with characteristics of invasive CRC

To investigate the origins of the NG2+ tumor subpopulation that we observed in the HCT116-derived VMT, we performed pseudotemporal reconstruction of differentiation trajectories using Monocle^32^. To interrogate the hierarchical relationships between the tumor subpopulations, HCT116-derived VMT scRNAseq data were subset in Seurat to contain only tumor A-D subpopulations by removing EC and fibroblast subpopulations. Applying Monocle to this subsampled population yielded a connected differentiation trajectory that separated into three main branches representing distinct cellular states – with tumor A and tumor C occupying two cell states and the NG2+ tumor subpopulation D occupying the third state along pseudotime (Figure 5A). Repeating this process with fibroblast subpopulations present in the subsampled dataset yielded a surprising finding: that fibroblasts segregate with tumor D to occupy the same state (Supplemental Figure 4). We next determined a list of genes that may be responsible for the transition of HCT116 CRC epithelial cells to a de-differentiated, mesenchymal-like state within the VMT (Figure 5B). We observed increases in EMT regulatory factors in the NG2+ tumor population, including TWIST1, VIM, TIMP1, and FN1, and decreased expression of differentiation and cellular adhesion markers, such as EPCAM, KRT8, KRT9 and KRT18 (Figure 5B). KRT5 and KRT23 serve as transitional markers for the tumor C population midway between tumor A and tumor D/fibroblast (mesenchymal) populations, with low expression at the endpoints of pseudotime and high expression at the midpoint. While KRT5 is associated with metastasis^29^, a recent study found that KRT23 activates CRC growth and migration, and that high KRT23 expression is prognostic of markedly shorter overall survival in CRC^33^. These findings suggest that HCT116 cells are undergoing EMT within the VMT, with each subpopulation occupying a distinct state. Indeed, using time-lapse microscopy we observed collective EMT events within the VMT, with rapid and coordinated movement of whole tumor clusters occurring within the span of an hour (Figure 5C, Supplemental Video 9). Interestingly, this is in contrast to the single-cell migratory phenotype we saw with HT29 cells reported above (Supplemental figure 1D).

**Figure 5:**
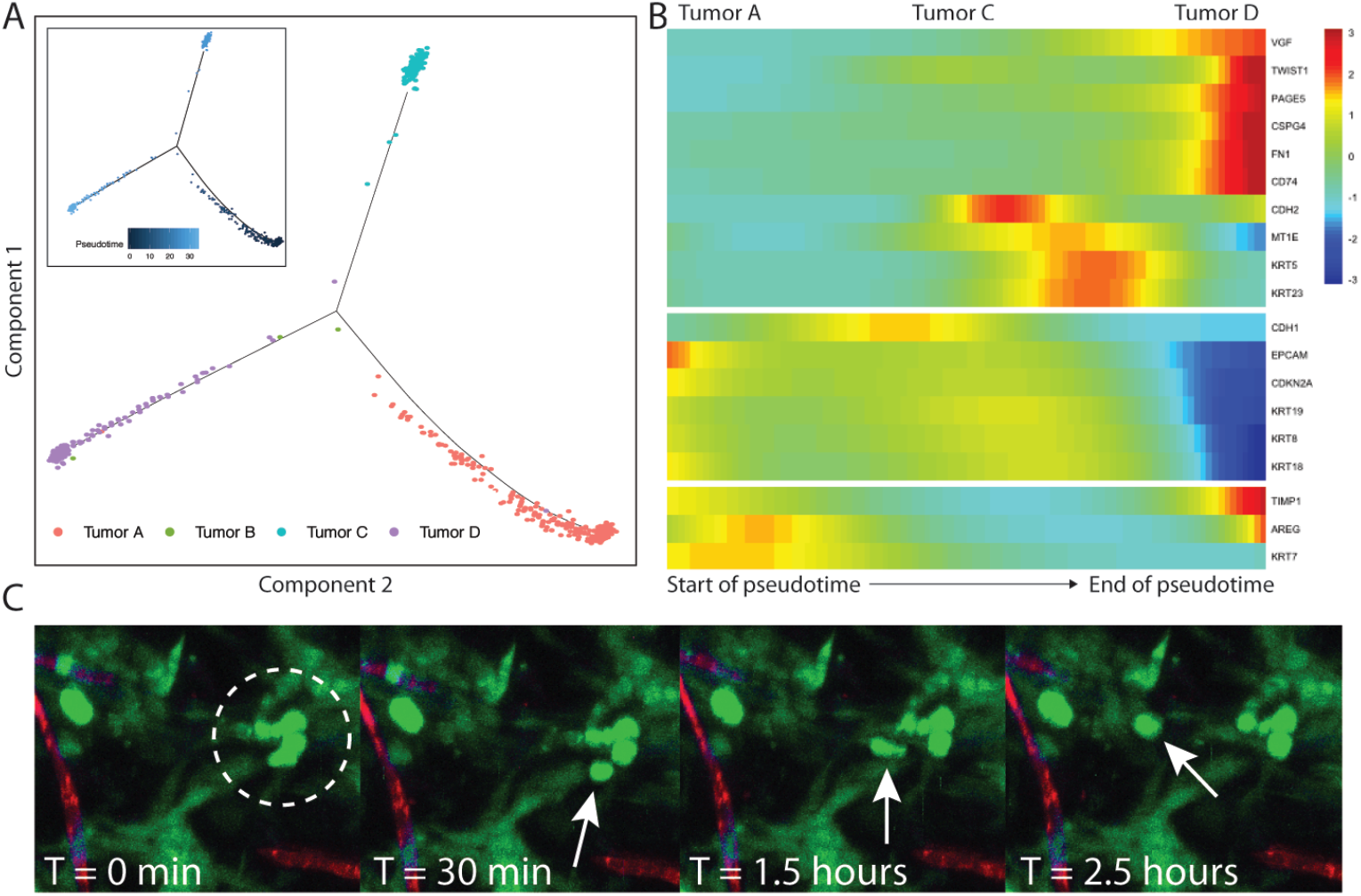
Lineage hierarchy reconstruction reveals differential tumor states between HCT116- and SW480-derived VMT. (A) Monocle analyses for HCT116-derived VMT tumor subset showing monocle trajectory plot and pseudotime plot (inset). (B) Pseudotime heatmap for HCT116-derived VMT transition subset showing 3 differentially-expressed clusters across the pseudotime trajectory. (C) Time-lapse confocal imaging showing an EMT event within the HCT116-derived VMT, characterized by collective and coordinated migration of tumor cells. Arrow shows movement of a cluster of tumor cells.

### SW480-derived VMTs retain *in vivo* tumor heterogeneity not seen in 2D or 3D monocultures

Given that SW480 CRC cells represent the major subtype of colorectal cancers (∼85%), we sought to validate the robustness of our system by performing additional scRNAseq on cells isolated from SW480-derived VMT, matched monolayer cultures, and tumor spheroids (Figure 6A). There is a growing use of tumor spheroids to model *in vivo* tumors as they capture the 3D nature of tumors and better model the cell-cell interactions. Similar to our findings with HCT116 we observed distinct shifts in SW480 gene expression between the VMT and monolayer cultures, and this was also seen for the EC and fibroblast populations (Supplemental Figure 5A,B). We also noted some overlap between VMT, monolayer and spheroid cultures for the SW480 tumor subpopulations. Populations were defined based on the same marker expression used for HCT116 (Supplemental Figure 5C), with a tSNE plot showing the discrimination between clusters (Supplemental Figure 5D). A heat map with the combined top 10 cluster-discriminative genes for SW480 populations showed that clusters were distinct in their gene signatures (Figure 6B). Similar to our findings with HCT116, two populations were identified as endothelial cells (EC A and EC B) by differential expression of PECAM1 (CD31+), three populations were derived from fibroblasts (fibroblast A, B and C) based on high expression of MMP2 and ACTA2, three populations were enriched for EPCAM and labeled as tumor (A, B and C), and a third tumor cluster highly expressed EPCAM and KRT5 (tumor D) (Figure 6B).

**Figure 6:**
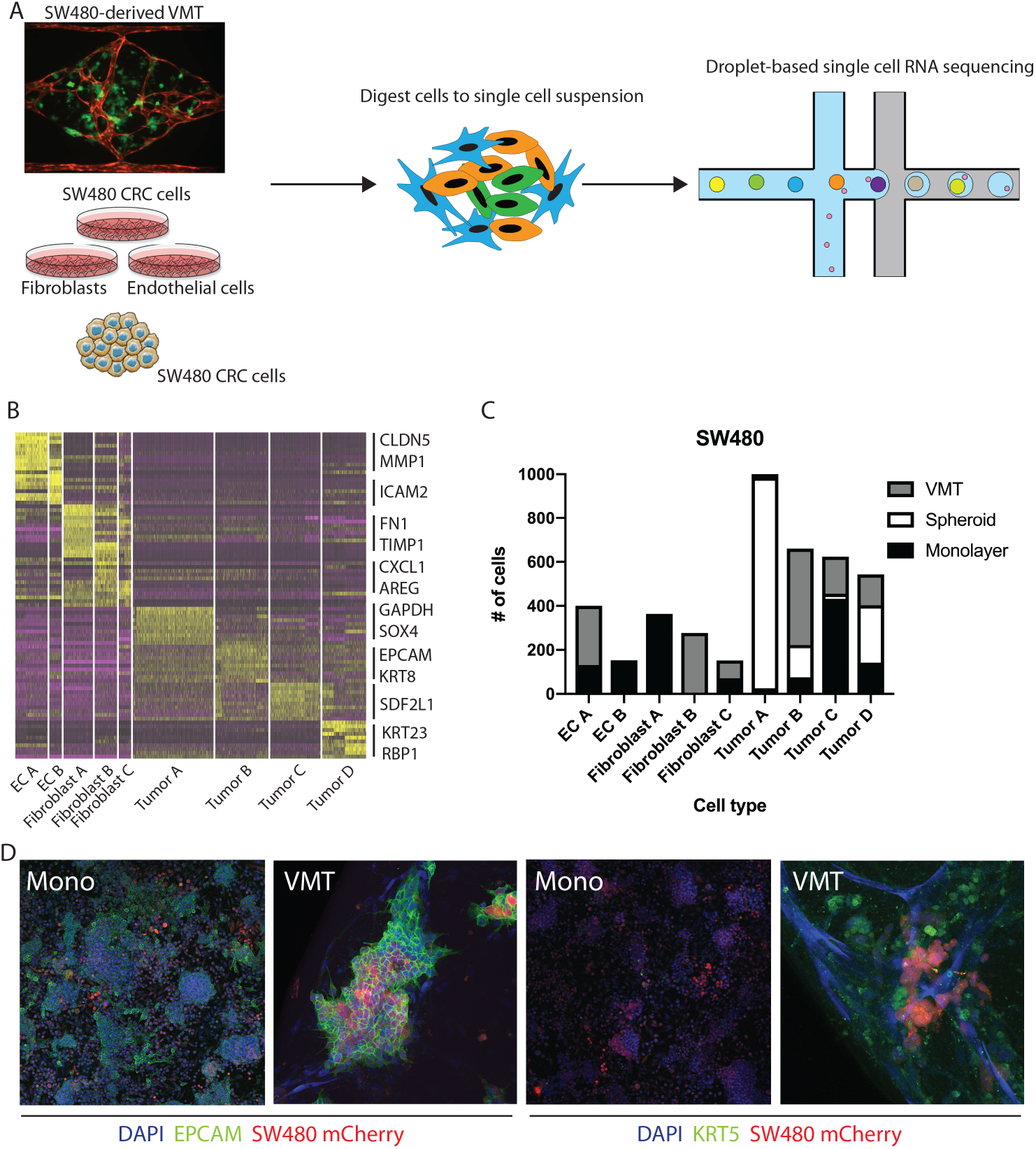
scRNAseq reveals SW480 tumor heterogeneity in the VMT platform. (A) Experimental design. SW480 CRC cells were grown as 2D or 3D (spheroid) monocultures, EC and fibroblasts were grown as 2D monocultures or all 3 cell types were co-cultured within the VMT, harvested on day 7 of culture and processed for droplet-based single cell RNA sequencing. (B) Heatmap of top 10 differentially expressed genes. (C) Distribution of cell types by sample. (D) Immunofluorescent staining of EPCAM and KRT5.

The distribution of each tumor type (A, B, C or D) between the different geometries was remarkably different, with tumor A dominating the spheroids, tumor B being enriched in VMT, tumor C dominating monolayer cultures, and tumor D being roughly evenly distributed between all geometries (Figure 6C). Similar skewed distributions were observed for the EC and fibroblast populations (Figure 6C). EPCAM and KRT8 characterized tumor B in the VMT, while tumor A, in the spheroids, had high levels of GAPDH and SOX4. Notably, the VMT had a more balanced distribution of tumor types than spheroids, which were dominated by tumor A. Interestingly, the NG2-enriched population seen in HCT116-derived VMT was absent from SW480. Similar to HCT116, however, we saw upregulation of EC PECAM1 in the VMT compared to monolayer cultures, and KRT5 expression in SW480 was enriched in the VMT compared to both 2D and 3D monocultures. EPCAM was not significantly enriched in a single group for the SW480 dataset (Supplemental Figure 5D,F). IF staining confirmed the presence of tumor populations expressing EPCAM or EPCAM/KRT5 in SW480 cells grown in monolayer and in the VMT (Figure 6D), with nearly ubiquitous expression of EPCAM in both systems, but higher expression of KRT5 in the VMT, confirming the scRNAseq data. We conclude that SW480 are more heterogeneous in the VMT geometry than they are in spheroids, which were dominated by a single type: tumor A.

### Pathways implicated in tumor progression are upregulated in CRC cells grown in the VMT when compared to cells grown as 2D or 3D monocultures

Comprehensive gene set enrichment analyses, based on cell type and state-specific marker genes, revealed pathways that were significantly upregulated in cells grown in the VMT compared to 2D or 3D monocultures (Figure 7A). Interestingly, the two tumor types show quite different patterns of gene expression, consistent with their different genotypes (SW480 is mutant for APC, KRAS and p53, while HCT116 is mutant for KRAS and PI3K but wild-type for p53 and APC). Both, however show a strong signal for glycolysis, ERBB1 and PDGF signaling, c-myc activation and integrin signaling. Consistent with the genotypes, we only saw up-regulation of β-catenin signaling in SW480 cells (target genes: JUN, XPO1, YWHAQ, ID2, CUL1, SP5, SFN; adjusted p-value 0.011). Within the VMT, tumor populations with high expression of Wnt signaling effectors (HCT116 tumor A and SW480 tumor B) also showed the highest expression of stem-like markers (Figure 7B), in agreement with *in vivo* findings that suggest a critical role for Wnt signaling in CRC cancer stem cell maintenance^34^.

**Figure 7:**
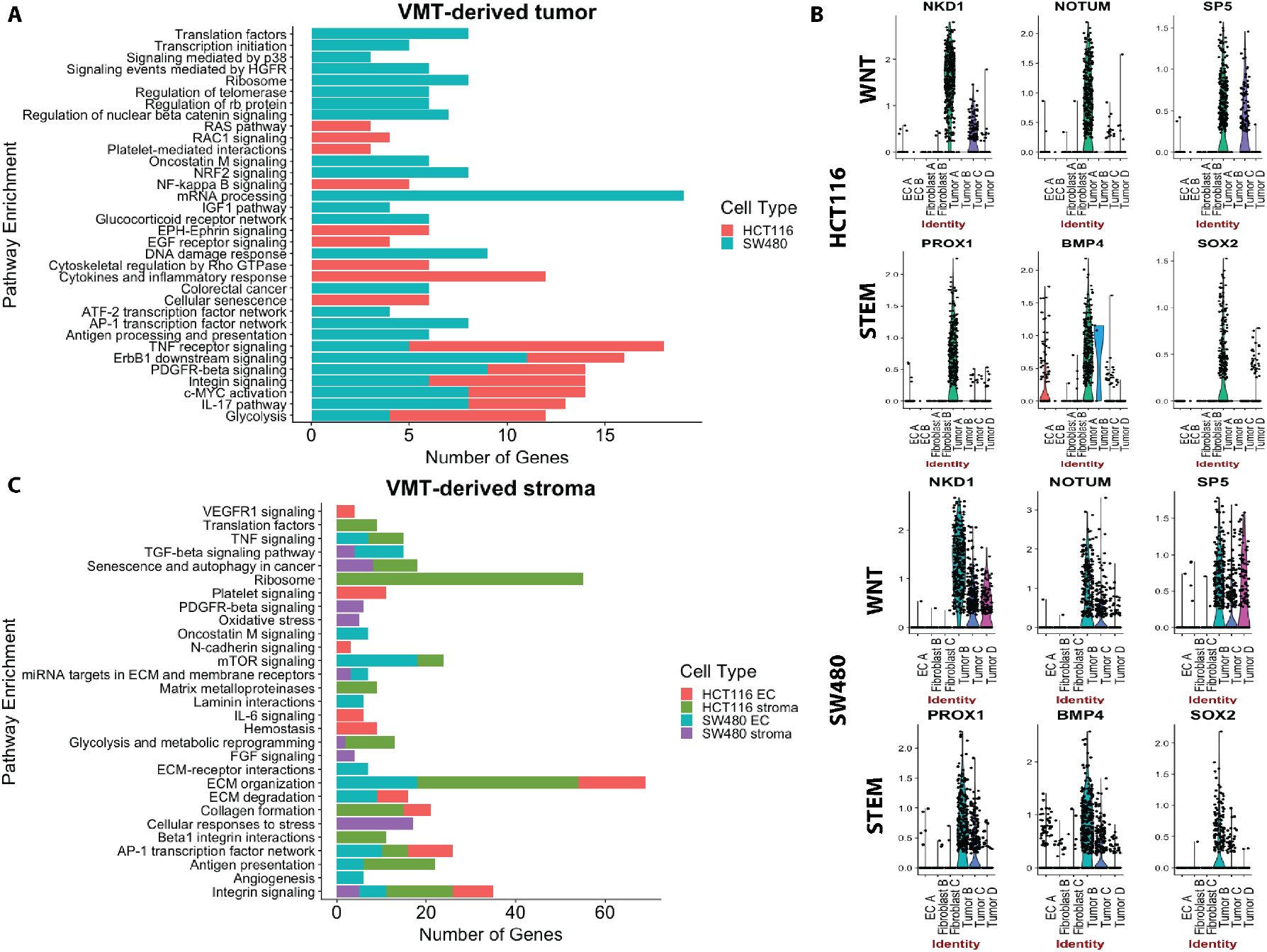
Pathway enrichment for VMT-derived cells. (A) VMT-derived tumor pathway enrichment for HCT116 and SW480. (B) Violin plots for Wnt-related genes and stem-like signatures for HCT116 and SW480. (C) Pathway enrichment for EC and stroma in HCT116 and SW480 VMTs.

When looking at the non-tumor cells in the VMT we noted a strong activation of the stroma by HCT116 versus SW480 tumor cells (Figure 7B), consistent with the more rapid and robust angiogenic response we see in the presence of HCT116 (unpublished observations). For example, HCT116 VMT-derived fibroblasts displayed upregulated uPA and uPA receptor-mediated signaling (ITGB1, LRP1, ITGB5, ITGA3, FN1; adjusted p-value 0.01) and increased expression of MMPs (MMP14, BSG, TIMP2, TIMP1; adjusted p-value 0.02). Furthermore, we observed an increase in β1 integrin cell surface interactions (ITGB1, CD81, COL4A1, LAMB2, ITGA3, COL6A2, COL6A1, FN1, COL6A3, LAMB1, LAMC1; adjusted p-value 0.0000017). EC within both HCT116- and SW480-VMT highly expressed genes associated with angiogenic sprouting and tip-cell formation, including ANGPT2, NID2 and DLL4, while expression of VWF, a marker of non-sprouting EC, was not differentially expressed between the VMT and monolayer cultures.

Compared to the SW480-derived VMT, spheroid cultures only showed upregulation of genes involved in HIF1α signaling and glycolysis, suggesting they are hypoxic; we found no other significantly enriched pathways, indicating that these 3D tumor spheroid models are limited in their ability to recapitulate features of *in vivo* tumors. In summary, we find that tumor-associated pathways are more highly activated in the VMT than in monolayer or spheroid cultures. Taken together, these findings suggest that the VMT, but not spheroid or monolayer culture, recapitulates a pro-tumorigenic microenvironment that promotes CRC pathogenesis.

### VMT-derived SW480 CRC cells better model *in vivo* tumor heterogeneity than 2D and 3D monocultures

To further validate the SW480-VMT as a physiologically relevant drug screening and disease modeling platform for mutant APC-driven CRC, we performed a reference-based integration using Seurat^35^. Cell anchoring removes technical variation between the separate scRNAseq experiments such that true biological variation can be detected above noise, and apart from batch effects. Of the three model systems, the VMT showed the greatest similarity to xenograft tumors, with 28% of tested cells accurately mapping to the reference dataset (Table 1). Monolayer and spheroid cultures did very poorly, with only 4% and 7% of cells, respectively, correctly mapping to the reference. Thus, even though these were not directly matched studies we still found a strong concordance between the VMT and *in vivo* tumors, confirming that the SW480-VMT is a superior model for capturing *in vivo* tumor complexity than are either monolayer or spheroid cultures.

**Table 1.** Results from reference-based integration of sw480 scRNAseq datasets

| Sample | # True | # False | % True | % False |
| --- | --- | --- | --- | --- |
| VMT | 188 | 486 | 28 | 72 |
| Spheroid | 50 | 624 | 7 | 93 |
| Monolayer | 24 | 650 | 4 | 96 |

### Targeting the tumor microenvironment suppresses tumor growth in the VMT platform but not in 2D or 3D monocultures

Interactions between tumor cells and their local environment are increasingly being seen as potential anti-tumor targets. Intriguingly, we observed significant upregulation of the TGF-β signaling pathway in fibroblasts derived from SW480-VMTs compared to those same fibroblasts grown in monolayers, while we saw no enhancement of TGF-β signaling in the SW480 cells themselves (Figure 8A). In contrast, HCT116 VMT-derived fibroblasts and tumor populations both showed modest increases in TGF-β signaling over 2D monocultures (Figure 8B). Importantly, we observed these differences in cancer-associated fibroblasts between SW480- and HCT116 VMTs despite using the same line and passage of fibroblasts (derived from a single vial) in all experiments, suggesting that each tumor line uniquely models the microenvironment and stromal compartment of the VMT. Indeed, these observations are consistent both with our scRNAseq data, and with recent literature defining the role of TGF-β-activated stroma in poor prognosis CRC^36^. Although most CRC display mutational inactivation of the TGF-β pathway, these tumors are paradoxically characterized by elevated TGF-β production. Functional studies show that cancer-associated fibroblasts support tumor-initiating cells via enhanced TGF-β signaling, and that poor clinical outcomes are predicted by a gene program induced by TGF-β in tumor-associated stromal cells^36^.

**Figure 8:**
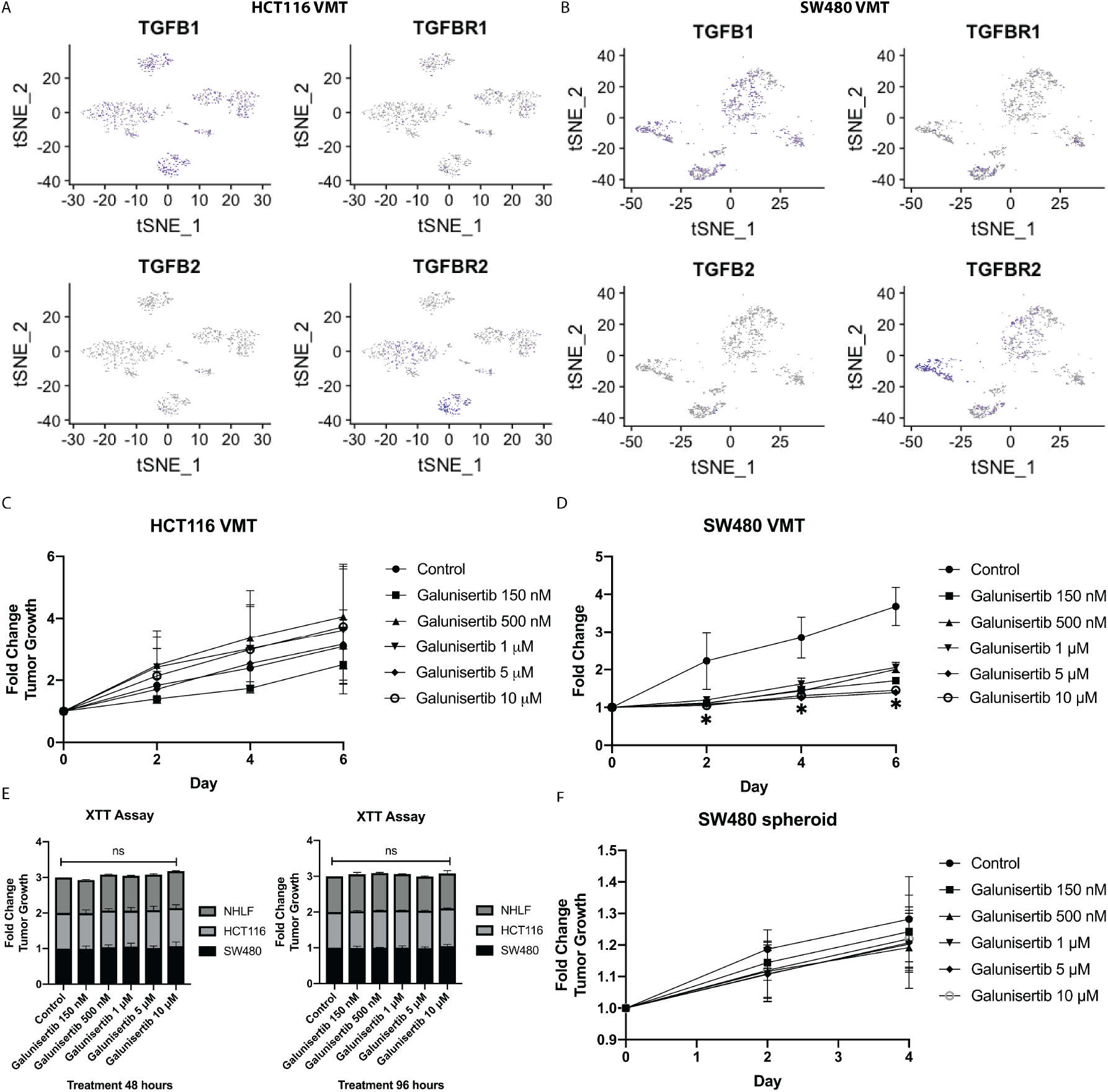
TGF-β-activated stroma in VMT confers differential drug sensitivity. (A) tSNE plot of HCT116 VMT with TGF-β ligands and receptors highlighted. (B) tSNE plot of SW480 VMT with TGF-β ligands and receptors highlighted. (C) Dose response of galunisertib for HCT116 VMT. (D) Dose response of galunisertib for SW480 VMT. (E) Dose response of galunisertib for monolayer HCT116, SW480 and NHLF at 48 hours and 96 hours. (F) Dose response of galunisertib for spheroid SW480.

Based on these findings we next tested whether targeting TGF-β signaling in the reactive stroma could suppress tumor growth in the VMT. We treated tumor cells in the VMT platform with the TGF-β receptor antagonist galunisertib, which is currently in clinical trials for hepatocellular cancer using a physiologically-relevant range of doses. We found that while galunisertib had no effect on HCT116 grown in the VMT over the course of 6 days (Figure 8C), it potently inhibited SW480 growth in the VMT (Figure 8D). Importantly, the drug had no effect on HCT116, SW480, or NHLF growing as 2D monocultures at any of the concentrations tested at 48 hours or 96 hours (Figure 8E). There were also no effects on SW480-derived spheroid growth in response to treatment (Figure 8F). These findings suggest that SW480 cells, but not HCT116, become dependent on TGF-β signaling in the presence of TGF-β activated stroma, and that the stroma is a powerful regulator of tumor growth as well as suppression. We conclude that the VMT model can serve as an important tool to further investigate molecular signatures of resistance or susceptibility to TGF-β antagonists, and could be used for tumor stratification and interrogation of additional tumorigenic pathways.

## Discussion

Anti-cancer drug development is a costly, time-consuming and inefficient process with a low success rate – only about 5% of drugs in the pipeline will ultimately gain FDA-approval. Recapitulating the complex intercellular crosstalk between the tumor and microenvironment is critical for increasing our understanding of clinical tumor pathology and laying the groundwork for novel therapeutic development. To this end, we show that the VMT more closely models the tumor growth and chemotherapeutic drug responses observed in preclinical *in vivo* murine xenograft models than 2D and 3D monocultures, with the added benefit of an entirely human cell system. In addition, the VMT captures tumor cell heterogeneity, vascular disruption and tumor-microenvironment interactions.

In the same way that malignancy progresses via aberrant signaling in the microenvironment, targeting key intercellular signaling pathways between cancer cells and stroma can abrogate malignant progression. High levels of TGF-β are associated with poor outcomes in CRC and a key hallmark of microsatellite-stable CRC (for which SW480 cells are a model) is TGF-β activated stroma. Furthermore, TGF-β can act as a tumor suppressor or tumor promoter depending on the context^37,38^. In VMT-derived stromal populations, we observed significant upregulation of TGF-β signaling that, when pharmacologically abrogated with galunisertib, resulted in suppressed SW480 tumor growth within the VMT, but had no effect on these cells in 2D or 3D monocultures, underscoring the critical role stromal cells play in disease progression and drug sensitivity. The VMT also captures the *in vivo* growth characteristics of tumors and response to standard chemotherapy that monolayer and spheroid cultures cannot. Both SW480 and HCT116 cells grow in the VMT at the same rate as they do *in vivo*, and in each case tumors show cessation of growth but not regression in response to FOLFOX. In sharp contrast, cell growth is rapid in monolayer and cell death is readily apparent after drug treatment. These findings highlight the unique capability of the VMT to recapitulate human-specific and clinically relevant tumor pathologies *in vitro* that cannot be readily reproduced with standard cell culture or animal models.

It is becoming increasingly clear that understanding cellular heterogeneity in tumors will be key to developing effective therapies. Our scRNA Seq studies reveal that the VMT captures this heterogeneity to a much higher degree than do either monolayer or spheroid cultures, and that only tumor cells in the VMT map strongly to tumor cells *in vivo*. Notably, the NG2+ tumor D population we see in HCT116 VMTs appears to arise through EMT, a phenomenon observed *in vivo* and within the VMT, but not in 2D or 3D monocultures. Thus, the VMT could be used to facilitate the study and targeting of resistance mechanisms related to this unique population, whereas this would not be possible with 2D or spheroid cultures, and would be difficult and possibly not human-relevant in mice.

Taken together, our findings demonstrate that the VMT not only recapitulates *in vivo* drug responses, but also reconstitutes the cellular diversity of the tumor growing *in vivo*. As such, we believe the VMT will prove a useful tool for drug discovery and drug validation, and may find utility in precision medicine settings.

## Methods

### Cell culture

Human endothelial colony-forming cell-derived endothelial cells (ECFC-EC) are isolated from cord blood with IRB approval. After selection for the CD31+ cell population, ECFC-EC are expanded on gelatin-coated flasks and cultured in EGM2 medium (Lonza). ECFC-EC are used between passages 4–8. Normal human lung fibroblasts (NHLF) are purchased from Lonza and used between passages 6–10. HCT116 colorectal cancer cells were donated from the UC Irvine Chao Family Comprehensive Cancer Center and SW480 colorectal cancer cells were clonally derived and provided by Dr. Marian Waterman. The ECFC-EC and cancer cells were transduced with lentivirus expressing mCherry (LeGO-C2, plasmid # 27339), green fluorescent protein (GFP) (LeGO-V2, plasmid # 27340), or azurite (pLV-Azurite, plasmid # 36086) (Addgene, Cambridge, Massachusetts)^39,40^. Cancer cells and fibroblasts are cultured in DMEM (Corning) containing 10% FBS (Gemini Bio). All cells are cultured at 37 °C, 20% O_2_, and 5% CO_2_.

### Microfluidic device fabrication and loading

Device fabrication and loading has been described previously^18,21^.

### Generation of tumor spheroids

SW480 CRC cells were resuspended into 8 mg/mL solution of 70% fibrinogen at a concentration of 1 × 10^5^ cells/mL and 50 µL of the cell solution was seeded into a 96-well plate containing 5 U of thrombin. Gels were allowed to set at 37°C for 15 minutes before addition of 100 µL of EGM2 medium to each well. Medium was changed every other day and drug treatment started on day 5.

### Drug treatment

Medium was changed every other day and hydrostatic pressure restored every day in the high throughput platform. After culturing for 5 days to allow full development of each VMT, culture medium was replaced by medium containing the drugs at the desired concentration, and delivered through the microfluidic channels using the hydrostatic pressure gradient. To mimic pharmacokinetics in the three *in vitro* systems (2D monoculture, 3D monoculture, and VMT), drugs were diluted into medium at maximal human plasma concentration on the first day of treatment and AUC were modeled according to average plasma clearance curves and drug half-life. Pharmaceutical grade 5-fluorouracil, leucovorin and oxaliplatin (FOLFOX) were purchased from UCI Medical Center pharmacy. Galunisertib (TGF-βR1 inhibitor) was purchased from SelleckChem. Each *in vitro* model system was exposed to compounds for the desired duration based on parameters defined by clinical administration, and the effect on tumor growth was quantified every 48 hours post-treatment. Cell viability in response to drugs in 2D monolayer cultures was quantified using an XTT assay according to the manufacturer’s protocol (Sigma-Aldrich). Briefly, 5,000 or 10,000 cells (HCT116, SW480 or NHLF) were seeded in triplicate in a 96-well plate and allowed to grow for 24 hours prior to treatment with drugs. XTT assays were performed after 48 hours of drug exposure or at the 96-hour time point after media changed at 48 hours. Cell viability was normalized to control wells without drug treatment.

### Immunofluorescence staining

VMTs were fixed for immunofluorescence staining by perfusing 4% paraformaldehyde (PFA) through the medium inlet for 30 minutes at room temperature. After fixing, VMTs were washed with 1x DPBS overnight at 4°C. Next, the high throughput plate was inverted and the bottom polymer membrane was carefully removed from the device. Each VMT unit was cut from the platform with a razor blade and placed in a 24-well culture plate, washed with 1x DPBS and then permeabilized for 15 minutes with 0.5% Triton-X100 diluted in DPBS. After permeabilization, VMT were blocked with 10% goat serum for 1 hour at room temperature and then incubated with primary antibody (diluted in 3% goat serum in 1x DPBS) overnight at 4°C. The following primary antibodies were used in experiments: anti-Collagen III (Abcam, ab6310), anti-EpCAM (Abcam, ab71916), and anti-Cytokeratin 5 (Abcam, ab53121). After washing with 1x DPBS, VMT were incubated with goat anti-rabbit or goat anti-mouse secondary antibody (1:2000 dilution in 5% serum) for 1 hour at room temperature before washing with DPBS and counter-staining with DAPI. Finally, anti-fade solution was added on top of each VMT before mounting with a glass coverslip. Established cell lines were seeded into multi-well cover glass chambers and processed by the same protocol.

### Fluorescence imaging and analyses

Fluorescence images were acquired with an Olympus IX70 inverted microscope using SPOT software (SPOT Imaging, Sterling Heights, Michigan). Confocal time-lapse series acquisition and imaging of fluorescent immunostaining was performed on a Leica TCS SP8 confocal microscope using a standard 10x air objective or 20x multi-immersion objective with digital zoom setting. AngioTool software (National Cancer Institute) was used to quantify vessel area, vessel length, number of vascular junctions and endpoints in the VMT. ImageJ software (National Institutes of Health) was utilized to measure vessel diameter and measure the total fluorescence intensity (i.e. mean grey value) for each tumor image to quantify tumor growth. Each chamber was normalized to baseline. Vessels were perfused by adding 25 µg/mL FITC- or rhodamine-conjugated 70 kDa dextran to the medium inlet. Once the fluorescent dextran had reached the vascular network, time-lapse image sequences were acquired using a Nikon Ti-E Eclipse epifluorescence microscope with a 4x Plan Apochromat Lambda objective. Perfusion images were analyzed using ImageJ software by measuring change in fluorescence intensity within regions of the extracellular space. Permeability coefficient is calculated by using Equation 1:

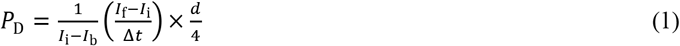

where *I*_*i*_, *I*_*f*_ and *I*_*b*_ represent the initial, final and background average intensities, respectively, ∆t is the time interval between two captured images and d is the average diameter of the vessel^41^. Any adjustments made to images are performed on the entire image, with all images in that experimental group adjusted to the same settings.

### Finite element simulation

COMSOL Multiphysics®5.2a was used to perform finite element simulations for the interstitial flow inside a developed microvascular network. Thresholded vessel images were smoothed and processed into outlines using ImageJ software, then converted into a .dxf file using Img2cad software. After the vessel outline was closed, and redundant fragments were removed using AutoCAD software, the complete vessel outline was scaled and integrated into the geometry of a microfluidic device. The refined CAD vessel diagram was then built into a 2D free and porous media flow model in COMSOL Multiphysics. Water was chosen to model the flow of culture media through the vascular network. The porosity and permeability of fibrin gel were estimated to be 0.99 and 1.5 × 10^−13^ m^2^, respectively, based on our published result^18^. Inlet/outlet were designated at the media reservoir boundaries, with pressure specified as 98 and 0.001 Pa, respectively, based on calculated gravity-driven pressure difference in the device, as previously described^18^.

### Cell sorting

The PDMS membrane was carefully removed from the bottom of the platform to expose the VMT tissue chambers. Chambers were washed with HBSS before adding TrypLE express enzyme drop wise to each VMT unit and incubated at 37°C for 5 minutes to loosen the extra-cellular matrix from the device. Chambers were flushed with a pipette to collect cells into a conical tube and washed once with EGM2 medium to stop the digestion reaction. Next, cells were subjected to centrifugation (340×g for 3 min.) and briefly resuspended in an HBSS/collagenase type III solution (1 mg/mL) to fully digest the matrix. Finally, the single cell suspension was washed with EGM2 and put through a 70 µm filter. Cells were processed for sorting within an hour and kept on ice. Sorting was performed on the BD flow assisted cell sorting (FACS) AriaII with BD FACS Diva software version 8.0.1. Samples were sorted using a 100 µm nozzle at 20 psi with a gating strategy to select for fluorescently labeled (mCherry) cancer cells (Supplemental Figure 6). Note that mCherry labeled CRC cells are pseudocolored green in all figures, while EC (vessels) are colored red.

### Animal studies

All animal experiments were approved by the University of California, Irvine (UCI) Institutional Animal Care and Usage Committee (IACUC). For CRC xenograft treatment experiments, male NOD SCID gamma (NSG) mice (Jackson Laboratories) were injected subcutaneously into both flanks with a sterile culture of HCT116 or SW480 (5 × 10^5^ cells) in 100 µL PBS (2 injections per mouse). Tumor volume and body weight were measured every other day using a caliper and scale, respectively. Tumor volume was calculated using Equation 2 with length being the longest measurement of the tumor.

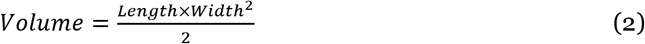

When the tumors reached a volume of 150 mm^3^, mice in the control group received two intraperitoneal (I.P.) injections of PBS. Mice in the FOLFOX group received one I.P. injection containing a combination of leucovorin (90 mg/kg) and 5-Fluorouracil (50 mg/kg), followed by an injection of oxaliplatin (6 mg/kg) two hours later. This treatment occurred weekly for up to 6 weeks or until the tumor reached a volume of 2 cm^3^. For NanoString experiments, xenograft tumors were established as described. When the tumors reached a volume of 2 cm^3^, mice were euthanized and tumors were harvested. The tumor tissue was minced into 1 mm pieces and digested in a solution of collagenase IV, hyaluronidase and DNAse I by shaking for 1–2 hours at room temperature. The single cell suspension was then processed and subjected to flow cytometry to select for mCherry labeled cancer cells as outlined above.

### NanoString PanCancer Human Pathways Assay

Each biological replicate was established in parallel from a single vial and passage of HCT116 cells. Gene expression analysis was performed using the NanoString PanCancer Human Pathways Assay. Per sample, 10,000 cells were lysed in diluted Buffer RLT (1:5 in RNAse-free water) at a concentration of 2000 cells/µL. Samples were submitted to the Genomics High Throughput Core Facility at UCI. Each cell lysate was mixed with a 3’ biotinylated capture probe and a 5’ reporter probe tagged with a fluorescent barcode from the custom gene expression code set. Probes and target transcripts were hybridized at 65°C for 12–16 h. Hybridized samples were run on the NanoString nCounter preparation station using the recommended manufacturer protocol, in which excess capture and reporter probes were removed and transcript-specific ternary complexes were immobilized on a streptavidin-coated cartridge. The samples were scanned at maximum resolution on an nCounter Digital Analyzer. Data were processed and analyzed using NanoString nCounter nSolver software version 3.0 (NanoString Technologies, Seattle WA). For gene expression analysis, data were normalized using the geometric mean of housekeeping genes selected by the GeNorm algorithm. The raw count from NanoString was subjected to background subtraction and reference gene normalization.

### Xenograft tumors for single-cell sequencing and cell anchoring

SW480 CRC cells were transduced with lentivirus carrying pCDH vector from System Biosciences: empty vector (Mock) or vector expressing dnLEF-1, followed by selection with 500 µg/mL G418. Transduced cells were collected as a pool for confirmation of expression, and Wnt signaling activity was measured by a SuperTOPFlash luciferase reporter^42^. Next, 2.5 × 10^6^ cells in PBS were injected subcutaneously into immunodeficient NSG mice. Tumors were removed after 3 weeks and dissociated for single cell sequencing. Only empty vector (Mock) samples were used for reference integration.

### SW480 xenograft dissociation for single cell sequencing

Tumor digestion has been described previously^43^. Single tumor cells were resuspended to 5 × 10^7^ cell/mL and stained with CD298-APC (10 µL/10^6^ cells)(BioLegend) and Sytox Green (ThermoFisher) for 20 minutes at 4°C in the dark, prior to sorting on FACSAria II (BD) for Sytox-cells. Sorted cells were spun down and resuspended to 1 × 10^6^ cells per mL.

### Single-cell sequencing

Flow cytometry sorted cells were washed in PBS with 0.04% BSA and re-suspended at a concentration of ~1000 cell/µL. Cellular suspensions were loaded onto a Chromium Single Cell Instrument (10X Genomics) to generate single-cell gel beads in emulsion (GEMs). GEMs were processed to generate cDNA libraries by using 10X Genomics v2 chemistry according to the Chromium Single Cell 3’ Reagents Kits v2 User Guide: CG00052 Rev B. Quantification of cDNA libraries was performed using Qubit dsDNA HS Assay Kit (Life Technologies Q32851), high-sensitivity DNA chips (Agilent 5067-4626) and KAPA qPCR (Kapa Biosystems KK4824). Libraries were sequenced on an Illumina HiSeq4000 to achieve an average of 50,000 reads per cell.

### Transcriptome alignment and data processing

After demultiplexing sequencing libraries to individual library FASTQ files, each library was aligned to an indexed GRCh38 reference genome using Cell Ranger (10X Genomics) Count 2.2.0. Aligned libraries are then normalized based on median reads per cell using Cell Ranger Aggr 2.2.0. Raw gene expression matrices were loaded into R (version 3.6.0)^44^ and converted to a Seurat object using the Seurat R package (version 2.3.4)^27^. From this, the data were log-transformed and corrected for unwanted sources of variation using the ScaleData function in Seurat. Gene expression matrices were then normalized to total cellular read count and to mitochondrial read count using linear regression as implemented in Seurat’s RegressOut function. For quality control filtering, we excluded cells with less than 200 or more than 5000 genes detected, and over 15% UMIs (unique molecular identifiers) derived from mitochondrial genome. In addition, genes that were not detected in at least 3 cells after this trimming were also removed from further analysis. Cell cycle gene regression did not significantly influence PCs or clustering (Supplemental Figure 7) and therefore was not performed in the final analyses.

### Single cell genotyping analyses

In order to identify cell origin, Vartrix^45^ combined with in-house scripts were used to extract single cell variant information from 10X genomics single cell data for the two HCT116 samples, VMT and monolayer. Variant information for HCT116 cell line was obtained from the Broad Cancer Cell Line Encyclopedia^46^ containing 39,000 variants. The variant list was first filtered so that common variants in dbSNPs Build ID: 150^47^ were excluded. For the remaining 18K SNVs, Vartrix was used to extract genotype information at each locus from the BAM files of the two samples analyzed using cellRanger (10X Genomics). Further confirmation of these findings derived from additional barcoding based on Y-chromosome-linked genes since HCT116 CRC cells are derived from a male patient, and the fibroblast line used in the experiments were derived from a female patient (an EC origin for the cells was ruled out by a complete absence of any EC-specific markers). For each cell in the two samples, a genotype score was determined for all 18K SNVs by subtracting the expression of all X-linked genes to quantify the confidence that the cell is derived from HCT116. For copy number variation (CNV) analysis, we utilized a pipeline from the Broad Institute called inferCNV^48^. We combined all normal cells (endothelial cells and fibroblasts) as a non-malignant reference population and queried the unknown and malignant populations against this reference set using the following parameters: cutoff = 0.1, cluster by groups = False, HMM = False, and de-noise = True.

### Pathway analyses

Marker genes for each subgroup of cells were determined by Seurat following log-transformation. Based on cell type and state specific marker genes, comprehensive gene set enrichment was performed using Enrichr^49^. A p-value of 0.05 was used as a cut-off to determine significant enrichment of a pathway or annotated gene grouping.

### Cell anchoring

Cell anchoring was performed with Seurat version 3 using reference-based integration^35^. For these analyses, ‘Xeno’ (SW480 xenograft tumor scRNAseq dataset) was used as the ‘reference’, and all other SW480 datasets (VMT, spheroid and monolayer) were used as ‘query’. Briefly, canonical correlation analysis was performed followed by L2-normalization of the canonical correlation vectors to project the datasets into a subspace defined by shared correlation structure across datasets. Next, mutual nearest neighbors were identified across reference and query cells and served as ‘anchors’ to guide dataset integration. For each anchor pair, a score was assigned based on the consistency of the anchors across the neighborhood structure of each dataset. Plots were generated using Bioconductor package ‘projectR’^50^.

### Statistical analyses

Data are represented as mean ±standard error of at least three independent experiments, unless noted otherwise. Comparison between experimental groups of equal variance were analyzed using an unpaired t-test and 95% confidence interval or one-way ANOVA followed by Dunnett’s test for multiple comparisons. Statistical calculations were performed using GraphPad Prism 8.0. For Nanostring analyses, gene expression was averaged across each sample (4 replicates each) then the fold change of each sample (i.e. monolayer and VMT), using xenograft tumor as the reference group, was log2-transformed. The mean log fold change between monolayer and VMT, combining all genes vs each individual gene, was then compared using 2-sample t-test in SAS. Significance cut-off for all analyses was p < 0.05.

---

**Supplemental Figure 1:**
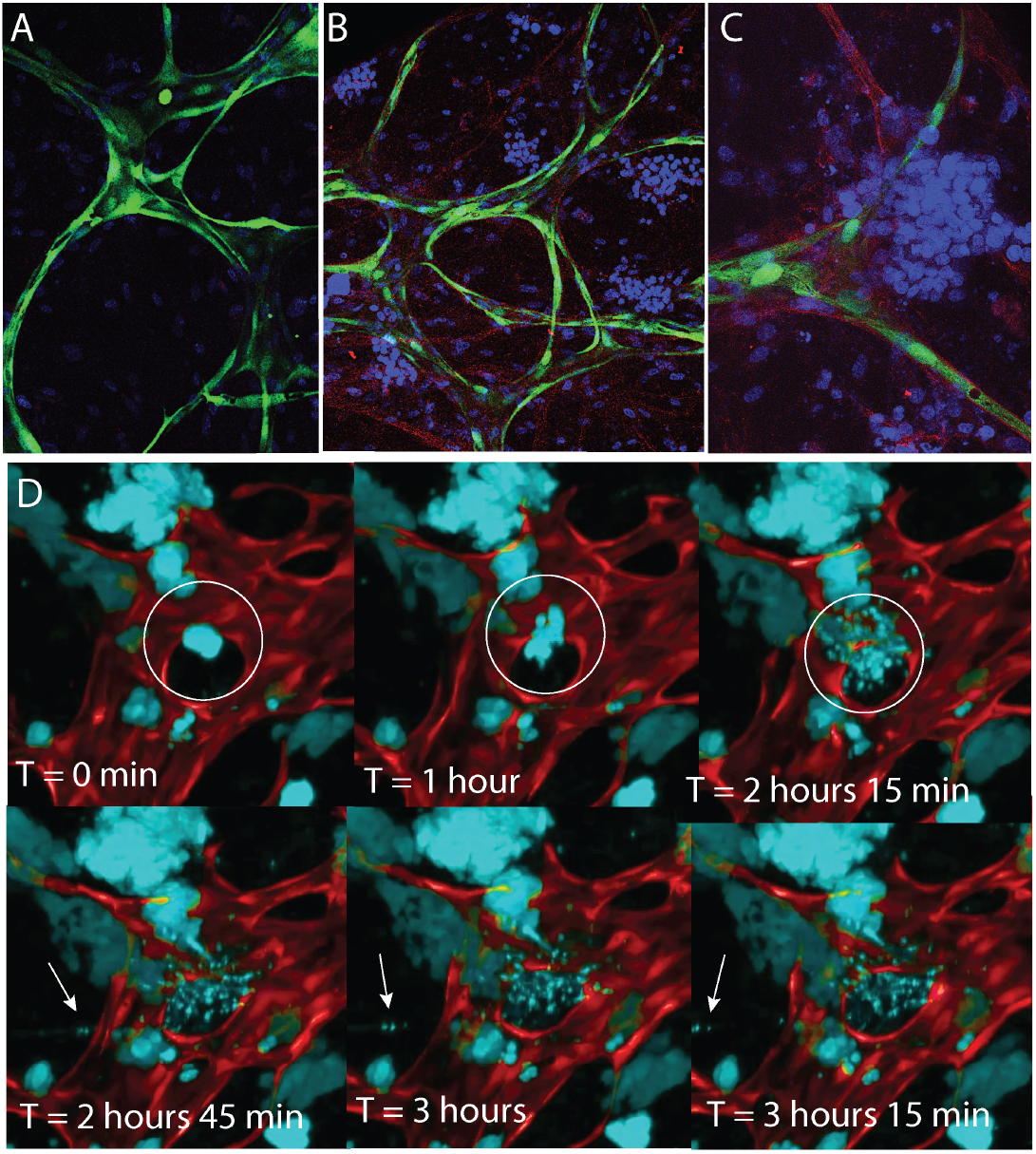
Collagen tracks revealed within the VMT. (A) Collagen III staining (red) is not apparent in the VMO. (B) Collagen III (red) is strongly expressed in the VMT. Cancer cells blue, vessels green. (C) Zoomed view reveals collagen fiber enrichment and ‘tracks’ in the VMT. (D) Time-lapse confocal imaging showing an EMT event within the VMT, characterized by collective and coordinated migration of tumor cells. Arrow shows movement of an individual CRC cell.

**Supplemental Figure 2:**
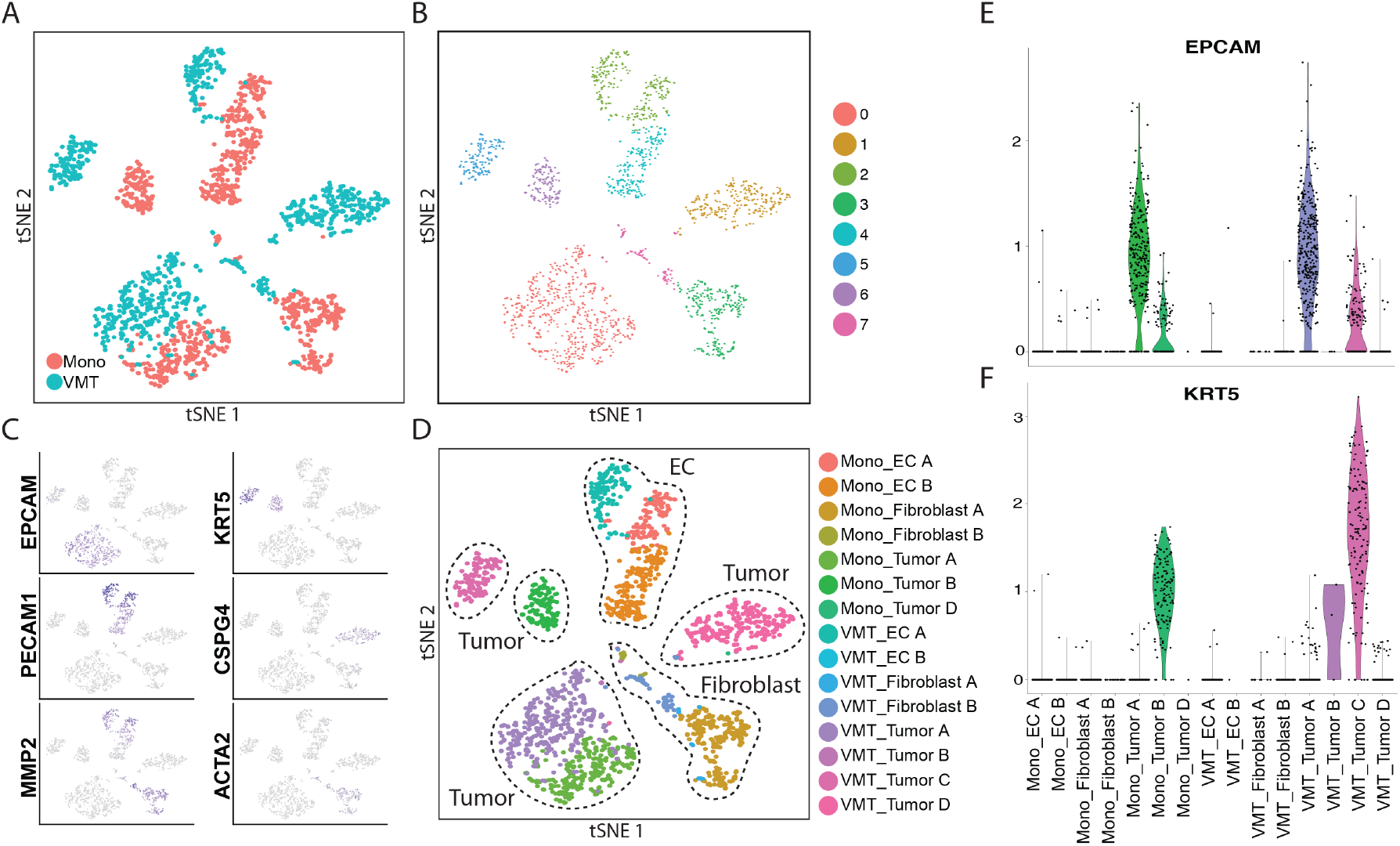
Single cell RNA sequencing reveals distinct changes between 2D monocultures and the VMT. (A) tSNE plot reveals marked shifts in gene expression between VMT and monolayer samples for all cell types. (B) Unbiased clustering for HCT116 dataset. (C) Cell clusters are characterized into types by known markers, and differential gene expression is displayed by tSNE plot. (D) tSNE plot shows each cluster annotated by group (Mono vs VMT) and cell type (EC A, EC B, Fibroblast A, Fibroblast B, Tumor A, Tumor B, Tumor C, Tumor D). Note that tumor C and D are absent from the monolayer culture. (E) Violin plot for EPCAM expression. (F) Violin plot for KRT5 expression.

**Supplemental Figure 3:**
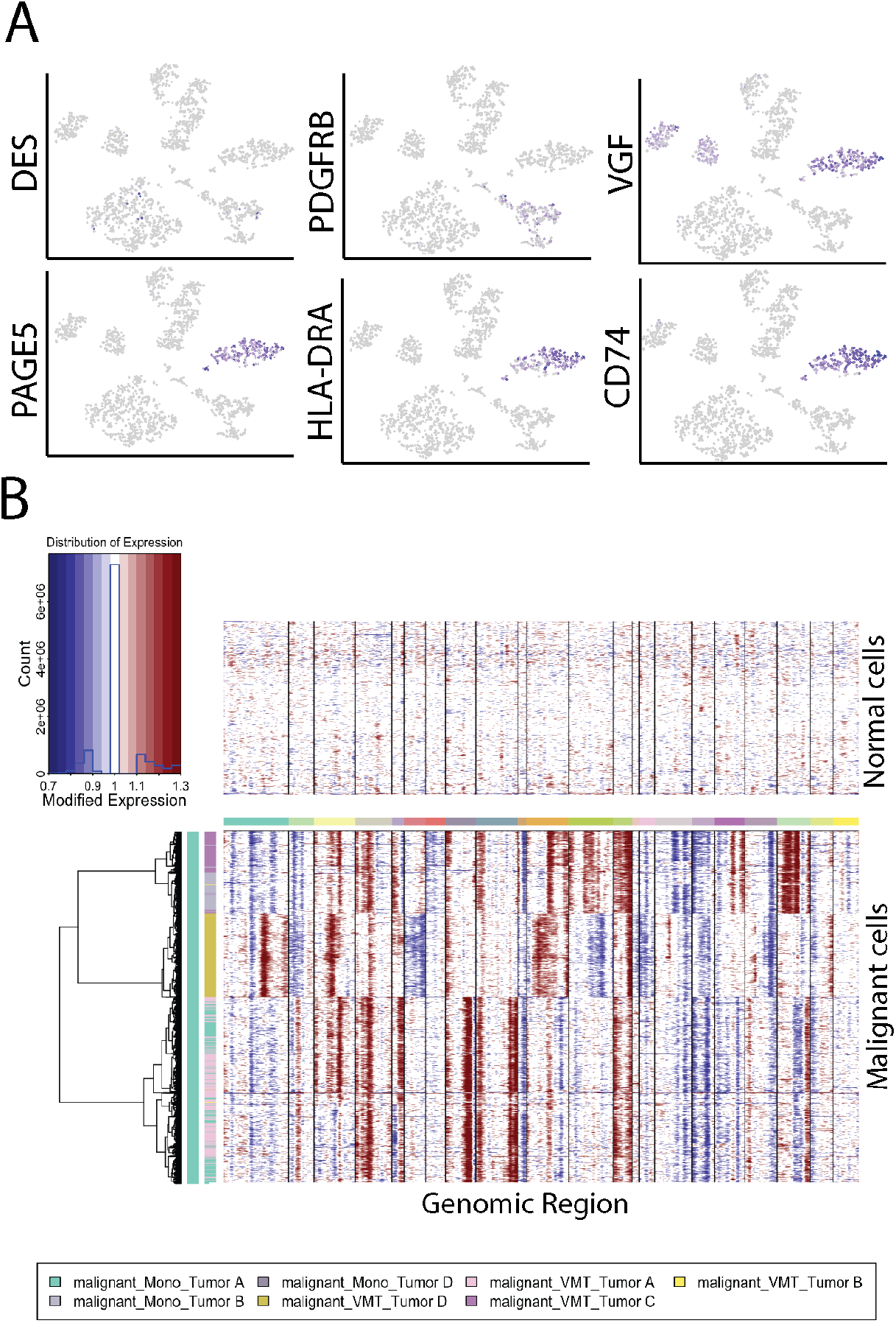
Single nucleotide variation and copy number variation analyses for HCT116. (A) tSNE plots showing marker gene expression for HCT116-derived VMT, highlighting the unique gene expression signature of tumor D. (B) CNV analyses showing that VMT-derived tumor D groups with known tumor-derived clusters A, B and C, whereas normal cells show greater chromosomal stability.

**Supplemental Figure 4:**
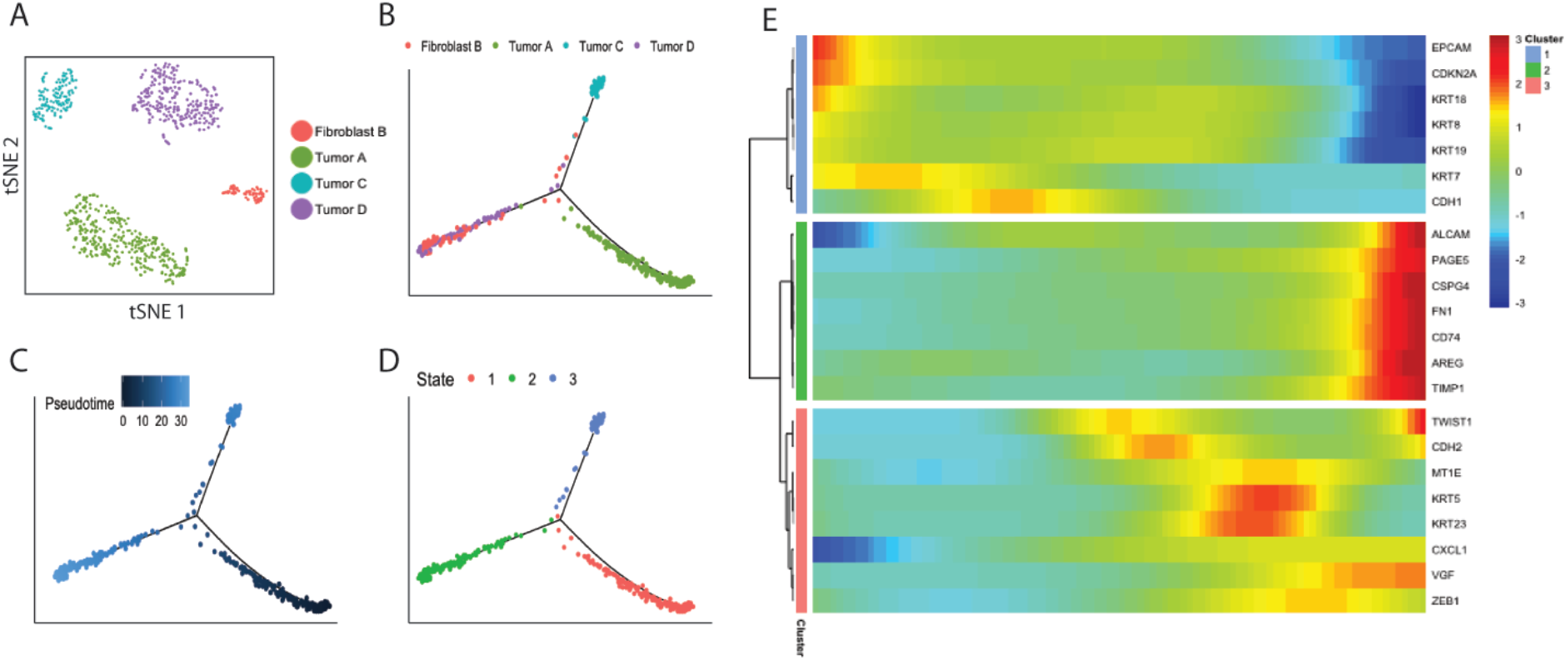
Differential tumor states observed in VMT. (A) Clustering for HCT116. (B) Monocle trajectory shows that tumor cells are distinct and occupy two distinct states, and that fibroblasts segregate to a third branch with tumor D to occupy the same state for HCT116. (C) Pseudotime plot of HCT116. (D) Cell state plot for HCT116. (E) Pseudotime heatmap for HCT116-derived VMT transition subset showing 3 differentially-expressed clusters across the pseudotime trajectory.

**Supplemental Figure 5:**
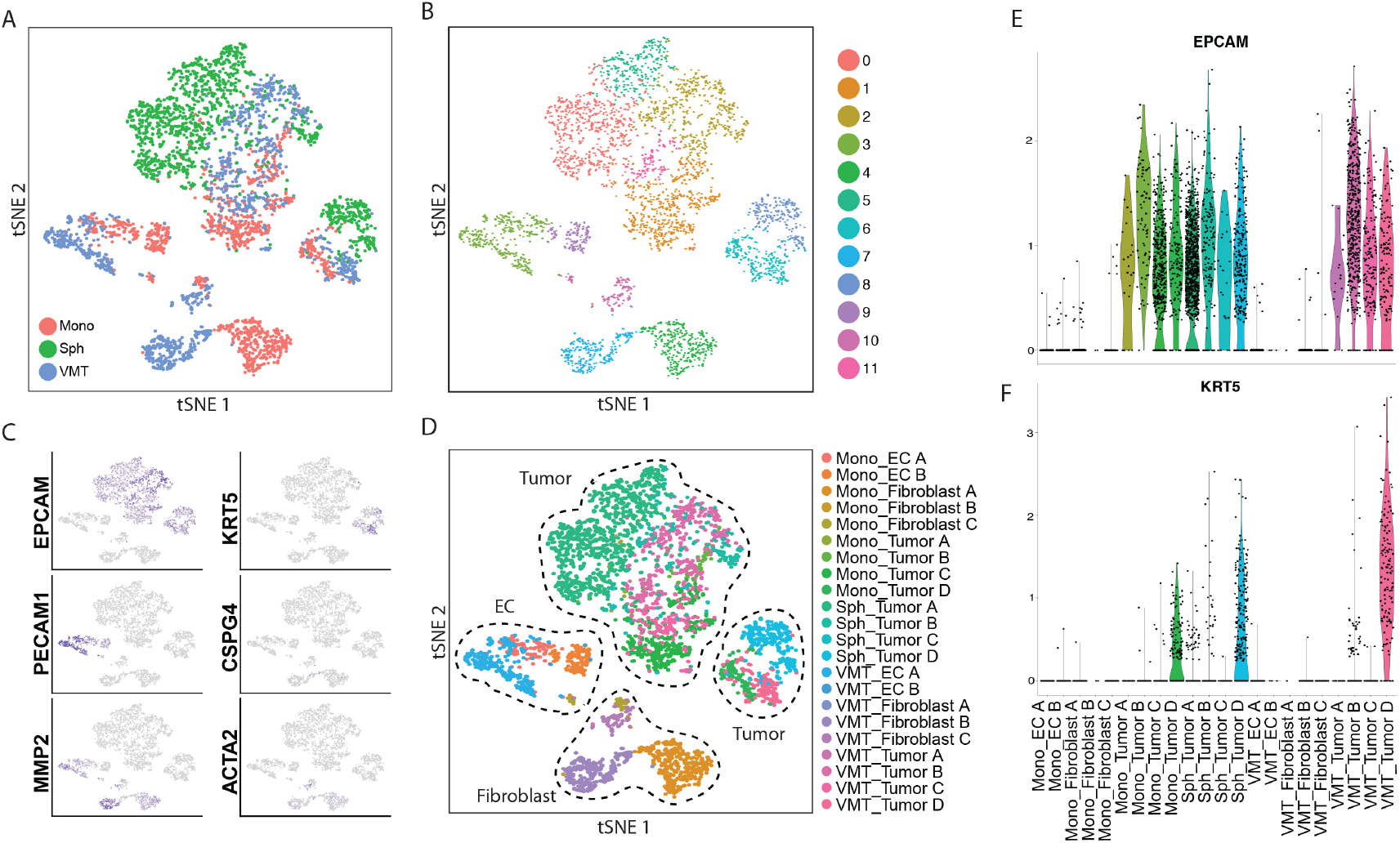
Single cell RNA sequencing reveals distinct differences between 2D monoculture, 3D monocultures (spheroids) and the VMT. (A) tSNE plot reveals marked shifts in gene expression between VMT, monolayer and spheroid samples for all cell types. (B) Unbiased clustering for SW480 dataset. (C) Cell clusters are characterized into types by known markers, and differential gene expression is displayed by tSNE plot. (D) tSNE plot shows each cluster annotated by group (Mono vs spheroid vs VMT) and cell type (EC A, EC B, Fibroblast A, Fibroblast B, Fibroblast C, Tumor A, Tumor B, Tumor C, Tumor D). Note that tumor C and D are absent from the monolayer culture (only 1 cell in tumor D). (E) Violin plot for EPCAM expression. (F) Violin plot for KRT5 expression.

**Supplemental Figure 6:**
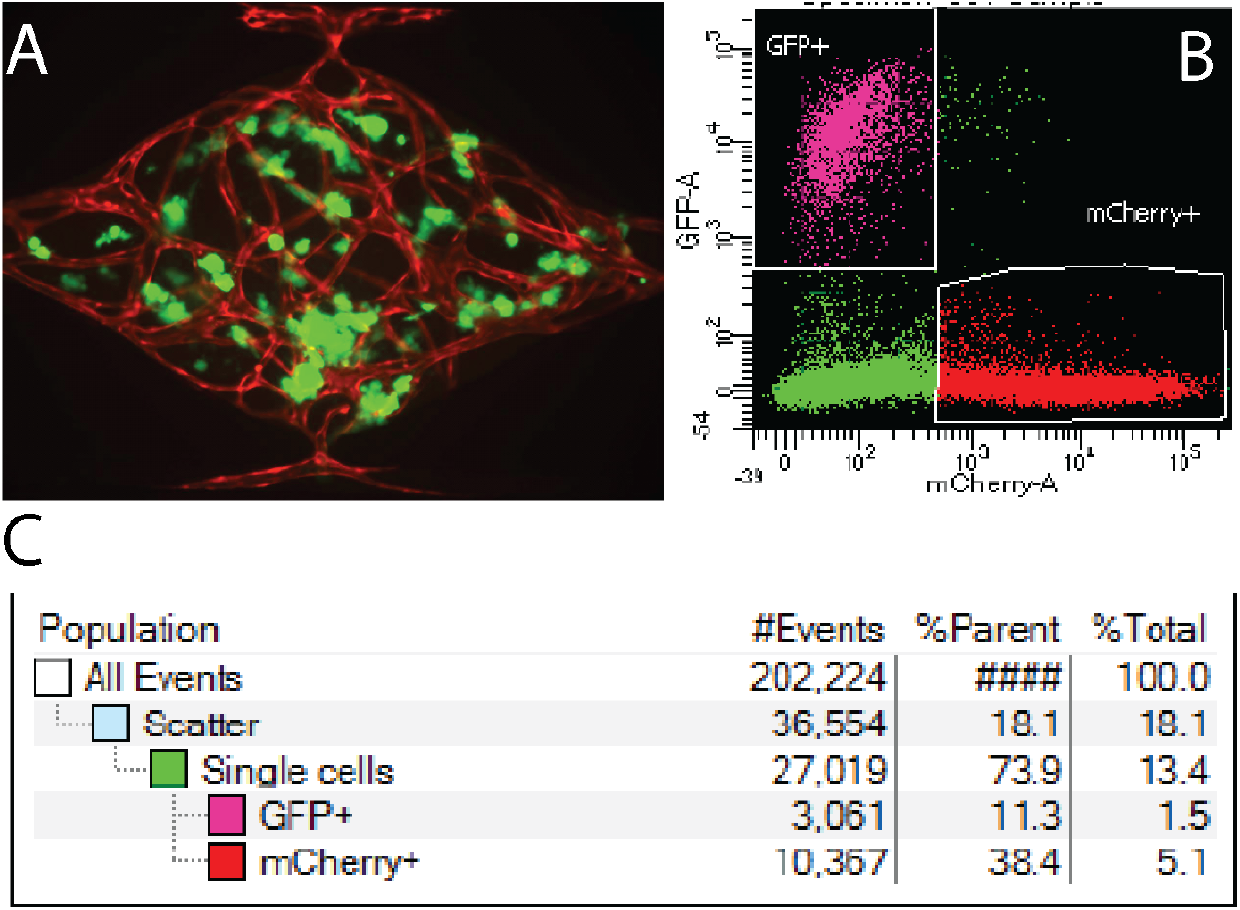
Cells in the VMT can be isolated via fluorescence activated cell sorting (FACS). (A) Representative image of SW480 VMT. CRC cells shown in green and vessels in red. Note that mCherry labeled CRC cells are pseudocolored green here for consistency with other figures. (B) Gating strategy for FACS. (C) Table showing proportions of isolated cells.

**Supplemental Figure 7:**
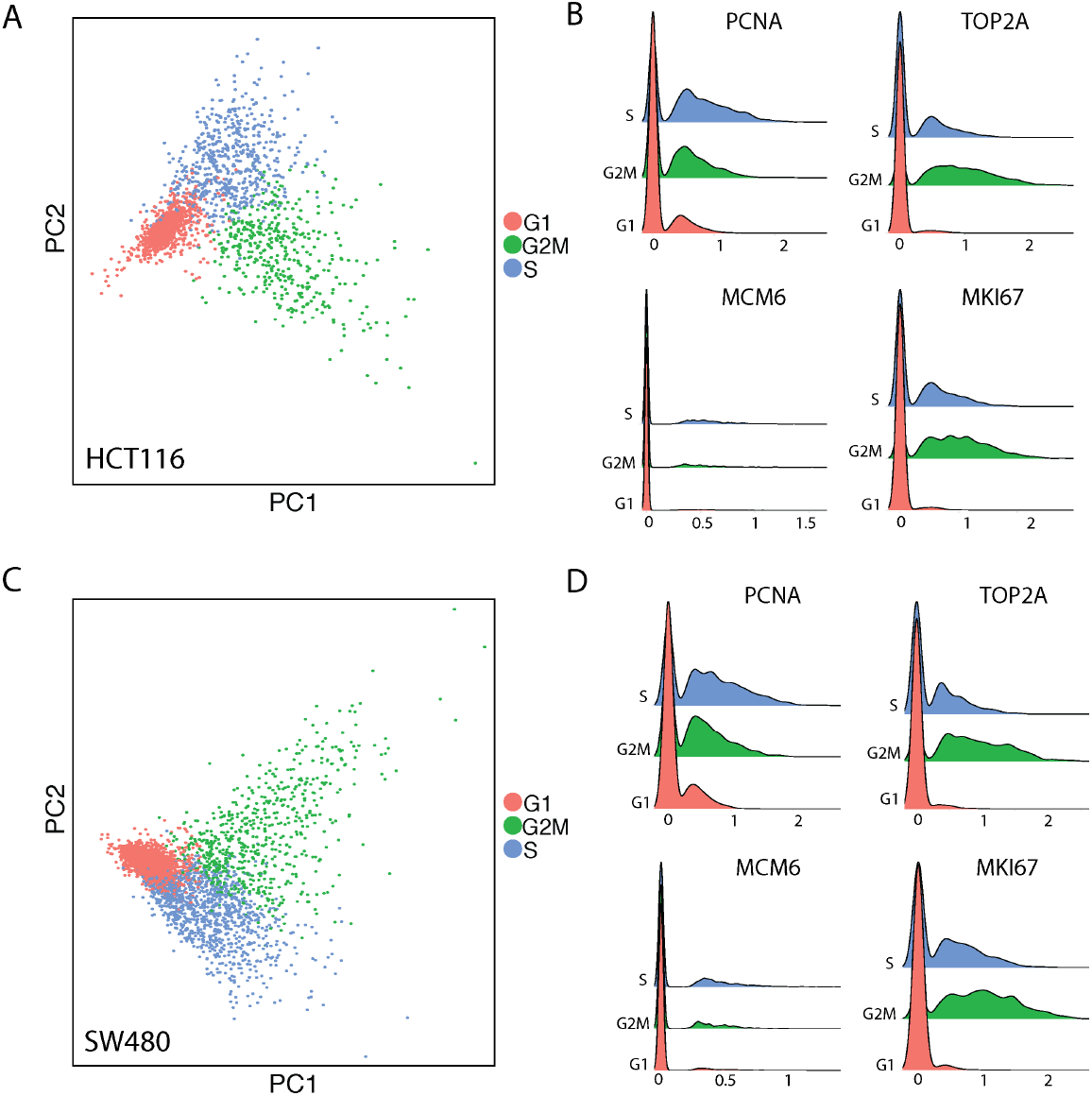
Cell cycle gene visualization for HCT116 and SW480. (A) Principal Component (PC) plot of HCT116 cells in G1, G2M, or S phase. (B) Representative cell cycle genes showing marginal upregulation across the HCT116 samples, with highest peak expression centered at 0 (no expression) and normal distribution with mean approximately 0.5 for expressing cell populations. (C) PC plot of SW480 cells in G1, G2M, or S phase. (D) Representative cell cycle genes showing marginal upregulation across the SW480 samples, with highest peak expression centered at 0 (no expression) and normal distribution with mean approximately 0.5 for expressing cell populations.

## Acknowledgements

We thank Damie Juat and Brittany Pham for help with cell culture, Hannah Bone for help with device fabrication, and Vanessa Scarfone for help with FACS experiments. S.J.H. is supported by an ARCS Award, Chancellor’s Club Fellowship, and Edwards Lifesciences Cardiovascular Technology Fellowship. This study has been supported by U54 CA217378, R01 CA180122, and UH3 TR-000481. C.C.W.H. and M.L.W. receive support from the Chao Family Comprehensive Cancer Center (CFCCC) through an NCI Center Grant [P30A062203]. The UCI Genomics High Throughput Facility and the Biostatistics Core are also supported by the CFCCC through this award.

## Author Contributions

S.J.H. performed the majority of the experiments and wrote most of the manuscript. S.M. performed the FOLFOX drug screening in the VMT and G.B.S. performed the FOLFOX and galunisertib drug screening in 2D and 3D monocultures. Q.H.N. performed scRNA-seq data preprocessing and helped with subsequent analyses and figure preparation. A.T. performed perfusion experiments comparing VMO and VMT. J.W. performed cell anchoring and cell genotyping analyses for the scRNA-seq datasets and wrote the corresponding methods. T.H. performed statistical analyses for the Nanostring dataset and wrote the corresponding methods. M.M.H. established xenograft tumors in mice. D.Z. performed COMSOL simulation, prepared the device figure and wrote the corresponding methods. E.C. and S.G. performed drug screening in the VMT. G.T.C. performed the xenograft scRNA-seq experiment. R.T.D. performed the CNV analyses and K.N. and N.P. helped with scRNA-seq analyses. D.A.L. and K.K. provided general guidance. A.P.L. and M.L.W. provided guidance and scientific feedback. C.C.W.H. directed the research and assisted in writing and editing the manuscript.

